# Ancient Ryukyu Jomon contributed to past and current genetic structure of Japanese populations

**DOI:** 10.64898/2026.04.03.712818

**Authors:** Masatoshi Matsunami, Yosuke Kawai, Leo Speidel, Kae Koganebuchi, Mai Takigami, Tsuneo Kakuda, Noboru Adachi, Yuichi Kameda, Chiaki Katagiri, Takayuki Shinzato, Akito Shinzato, Masami Takenaka, Naomi Doi, NCBN Controls WGS Consortium, Nancy Bird, Garrett Hellenthal, Minoru Yoneda, Takayuki Omori, Hiromasa Ozaki, Minoru Sakamoto, Naoko Kinoshita, Minako Imamura, Shiro Maeda, Ken-ichi Shinoda, Hideaki Kanzawa-Kiriyama, Ryosuke Kimura

**Author notes:** **Corresponding Authors:** Masatoshi Matsunami, PhD (lead contact), 1076, Kiyuna, Ginowan, Okinawa 901-2720, Japan, Ryosuke Kimura, PhD, 1076, Kiyuna, Ginowan, Okinawa 901-2720, Japan, Hideaki Kanzawa-Kiriyama, PhD, Amakubo 4-1-1, Tsukuba, Ibaraki 305-0005, Japan.

## Abstract

Characterized by the earliest use of pottery, the Jomon culture was a unique Neolithic culture that spread throughout the Japanese Archipelago. Previous archaeological evidence suggests that Jomon hunter-gatherers colonized the southernmost islands, the Ryukyu Archipelago, by approximately 7,000 years before present (YBP). However, genetic characteristics of the Ryukyu Jomon population and its contribution to the modern population have not been elucidated yet. In this study, we newly sequenced 273 modern and 25 ancient (6,700–900 YBP) whole genomes collected across the Ryukyu Archipelago. Our analysis demonstrated a genetic differentiation between the Hondo (Japanese mainland) and Ryukyu Jomon, dating back to ∼6,900 YBP. After the divergence from the Hondo Jomon, the Ryukyu Jomon experienced severe bottlenecks, with an effective population size of ∼2,000. Admixture between the Ryukyu Jomon and migrants from the historic Hondo population occurred ∼1,000 YBP, which corresponds to the widespread adoption of iron tools and agriculture in the Central Ryukyus. Different demographic histories between modern Hondo and Ryukyu populations resulted in different rates of Jomon ancestry in these populations. By providing a new perspective on the peopling of the Ryukyu Archipelago, this study significantly enhances our understanding of cultural transitions in the region.

## Introduction

The insular region of East Asia is one of the endpoints of the human journey. After the out-of-Africa migration approximately 50,000 years before present (YBP), anatomically modern humans spread across the world and reached the Japanese Archipelago by approximately 35,000 YBP (Fujita et al. 2016). The Japanese Archipelago stretches from north to south and is prehistorically divided into three regions: Hokkaido, the “Hondo” region (consisting of Honshu, Shikoku, and Kyushu), and the Ryukyu Archipelago.

The Ryukyu Archipelago, which is also called the Nansei Islands, constitutes the southernmost island chain of the Japanese Archipelago (Fig. 1A) and can be further divided into the Northern Ryukyus (the Osumi and Tokara Islands), the Central Ryukyus (the Amami and Okinawa Islands), and the Southern Ryukyus (the Miyako and Yaeyama Islands) based on archaeological and biological perspectives (Fig. 1B). Due to the historical and cultural differences between Hondo, the Central Ryukyus, and the Southern Ryukyus, the chronological period designations applied to these regions differ among them (Supplementary Fig. 1) (Pearson 2013).

**Fig. 1:**
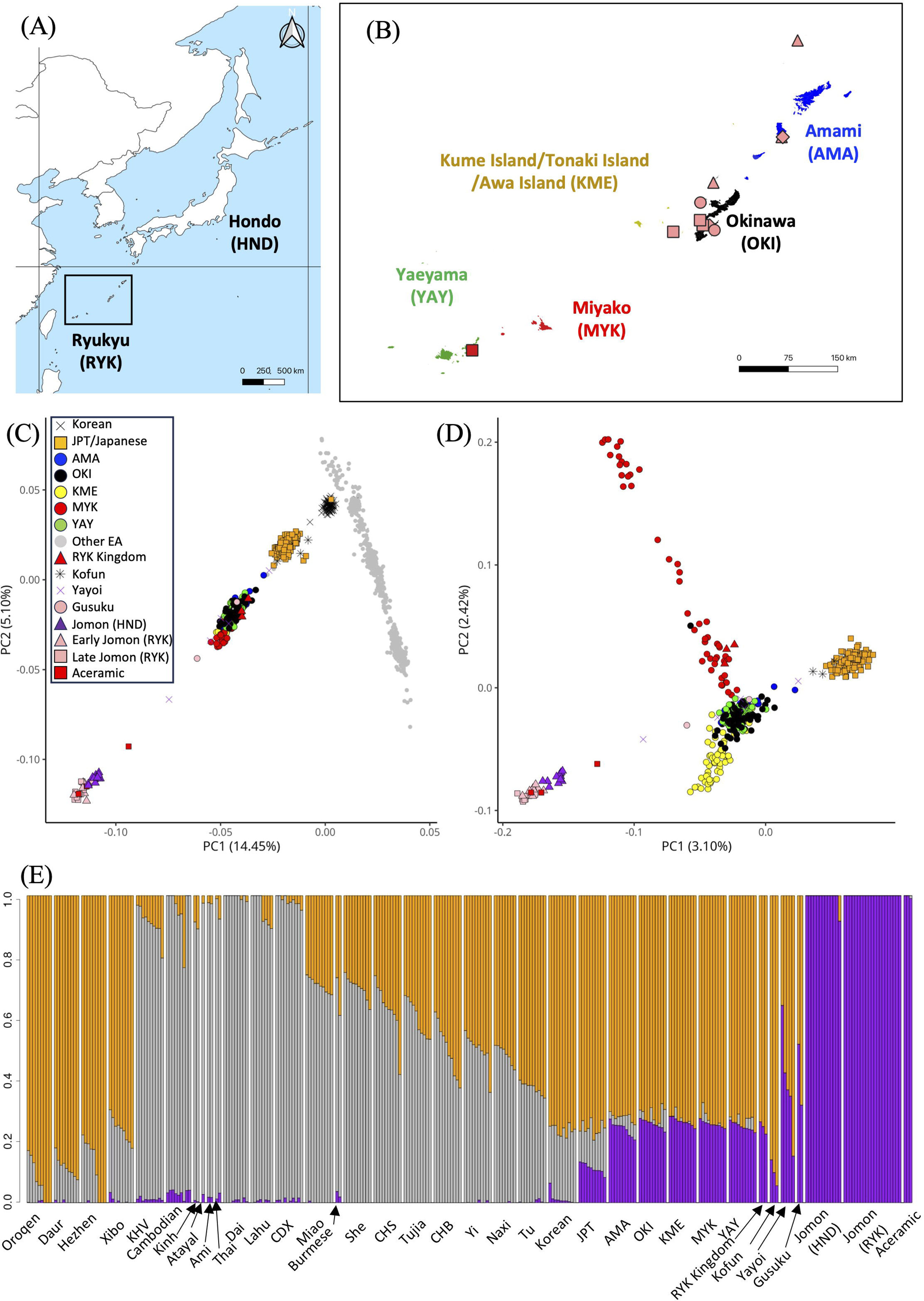
Integrated population genetic structure using modern and ancient Ryukyu genomes. **(A)** Geographic location of the Japanese Archipelago. The Japanese Archipelago can be divided into Hokkaido, Hondo (Honshu, Shikoku, and Kyushu Islands) and the Ryukyu Archipelago. **(B)** Geographic location and sampling points of ancient genomes in the Ryukyu Archipelago. The Central and Southern Ryukyu Archipelago includes Amami, Okinawa, Miyako, and Yaeyama Islands. The color of each island group and the shape of each sampling point of ancient genomes correspond to following PCA plot except for the Cliff bottom site at the Gushikawa Gusuku and the Tomachin site. Because these sites include both early and late Ryukyu Jomon, locations are represented by pink diamonds. **(C)** PCA plot using 1240 K SNPs from ancient and modern East Asian populations. Ancient genome data was projected into the modern genome plot. **(D)** PCA plot using only Japanese modern and ancient genomes. Coloring corresponds to **C**. **(E)** Admixture plot using K=3, which has the lowest cross validation errors. We randomly selected ten individuals from each modern population, when a population had data available for more than ten individuals. For ancient samples, we used samples showing a depth of more than 0.01.

In the Hondo region, although numerous Upper Paleolithic artifacts have been excavated, the only human bones dating back to the Paleolithic period are the Hamakita human remains (Kondo and Matsu’ura 2005). In contrast, there are many sites where Upper Paleolithic human bones have been excavated from the Ryukyus, including at the Minatogawa Fissure site (16,000 YBP) (Suzuki and Hanihara 1982) and the Shiraho-Saonetabaru Cave site (20,000 YBP) (Nakagawa et al. 2010) in the Central and the Southern Ryukyus, respectively. It is worth noting that microblade industries, which showed a common distribution in the Hokkaido and Hondo regions during the Upper Paleolithic period, were not present in the Central and Southern Ryukyus (Sato and Morisaki 2022). Therefore, the origins of these different Paleolithic cultures have been under debate.

After the Upper Paleolithic period, the Jomon pottery culture, developed by Neolithic hunter-gatherers, began around 16,000 YBP in the Hondo region (Habu 2004). By approximately 7,000 YBP, the pottery culture had been introduced to the Central Ryukyus (Takamiya et al. 2019; Pearson 2013; Fujita et al. 2016). This culture in the Central Ryukyus, known as the Shell-mound culture or the Ryukyu Jomon culture, continued even after the introduction of rice cultivation to the Hondo region during the Yayoi period (3,000 YBP), and ultimately lasted until the late Heian period (1,000 YBP). Therefore, the Neolithic culture in the Central Ryukyus can be divided into two periods: the Early Ryukyu Jomon period, which corresponds to the Jomon period in the Hondo region, and the Late Ryukyu Jomon period, which overlaps with the Yayoi to Heian periods.

In contrast, the Jomon culture did not extend to the Southern Ryukyus, resulting in a marked gap in the archaeological record. In the Yaeyama Islands and on Tarama Island, part of the Miyako Islands, the Shimotabaru culture (4,200–3,500 YBP) emerged, but subsequently disappeared. Later, the Aceramic culture (2,500–900 YBP), characterized by giant clam shell adzes, appeared on both the Miyako Islands and the Yaeyama Islands (Asato 1993; Yamagiwa et al. 2019). The origins of these two cultures remain a subject of debate. The cultures of the Central and Southern Ryukyus were finally unified during the Gusuku period (900–500 YBP), a time that saw the widespread adoption of iron tools and agriculture. This cultural transformation led to a period of warfare and eventually to the establishment of the Ryukyu Kingdom.

Recent genomic analyses have provided insights into human migration to the Ryukyu Archipelago. Previous studies on present-day individuals revealed clear genetic differentiation between the Hondo and Ryukyu regions and among island groups within the Ryukyu Archipelago (Yamaguchi-Kabata et al. 2008; Sato et al. 2014; Liu et al. 2023; Liu et al. 2024). Attempts have also been made to unravel the history of population formation in the Ryukyu Archipelago (Matsunami et al. 2021; Cooke et al. 2023; Koganebuchi et al. 2023; Yamamoto et al. 2024). For reconstruction of the population formation history, inclusion of ancient genomes makes the analysis more effective, but ancient genomic data from the Ryukyu Archipelago remain limited and fragmentary. To date, several ancient mitochondrial DNA (mtDNA) datasets, including those from the Upper Paleolithic period, have been reported (Shinoda et al. 2012; Shinoda et al. 2013a; Shinoda and Adachi 2017; Shinoda et al. 2018; Mizuno et al. 2021); however, ancient nuclear genome data are currently restricted to skeletal remains excavated from the Nagabaka site on Miyako Island, dating from the Aceramic to the Ryukyu Kingdom periods (Robbeets et al. 2021). Although it is notable that an individual from the Aceramic period had a genetic profile comparable to that observed in Jomon individuals, a comprehensive understanding of the population history in the Ryukyu Archipelago from ancient times to the present is still lacking.

In this study, we conducted whole-genome sequencing (WGS) of 273 modern and 25 ancient individuals (dating from 6,700 to 900 BP) from the Ryukyu Archipelago. Providing evidence for genetic differentiation between ancient populations in the Hondo and Ryukyu regions, we estimated the prehistoric population dynamics of the Ryukyu Archipelago.

## Results

### Dating and relatedness of sequenced ancient samples

In this study, we newly sequenced 25 ancient individuals from 12 archaeological sites across the Ryukyu Archipelago and directly confirmed the date of each bone sample (Fig 1B; Supplementary Fig. 2). Archeological evidence indicates that these samples are derived from the Ryukyu Jomon period and the Aceramic period in the Central and Southern Ryukyus, respectively (Supplementary Note 1). Radiocarbon dating results reported in previous studies for 19 of the 25 individuals, and the 6 newly analyzed individuals are shown in Supplementary Table 1. Although most of the radiocarbon dates for the ancient bones corresponded to the archeological estimation, two samples—from Gushibaru Shell Mounds and Tsuken Shell Mounds—were dated to the Gusuku period, which is later than the Ryukyu Jomon period (Table 1). Two human remains unearthed on Ishigaki Island were dated to the Aceramic period.

**Table 1.**
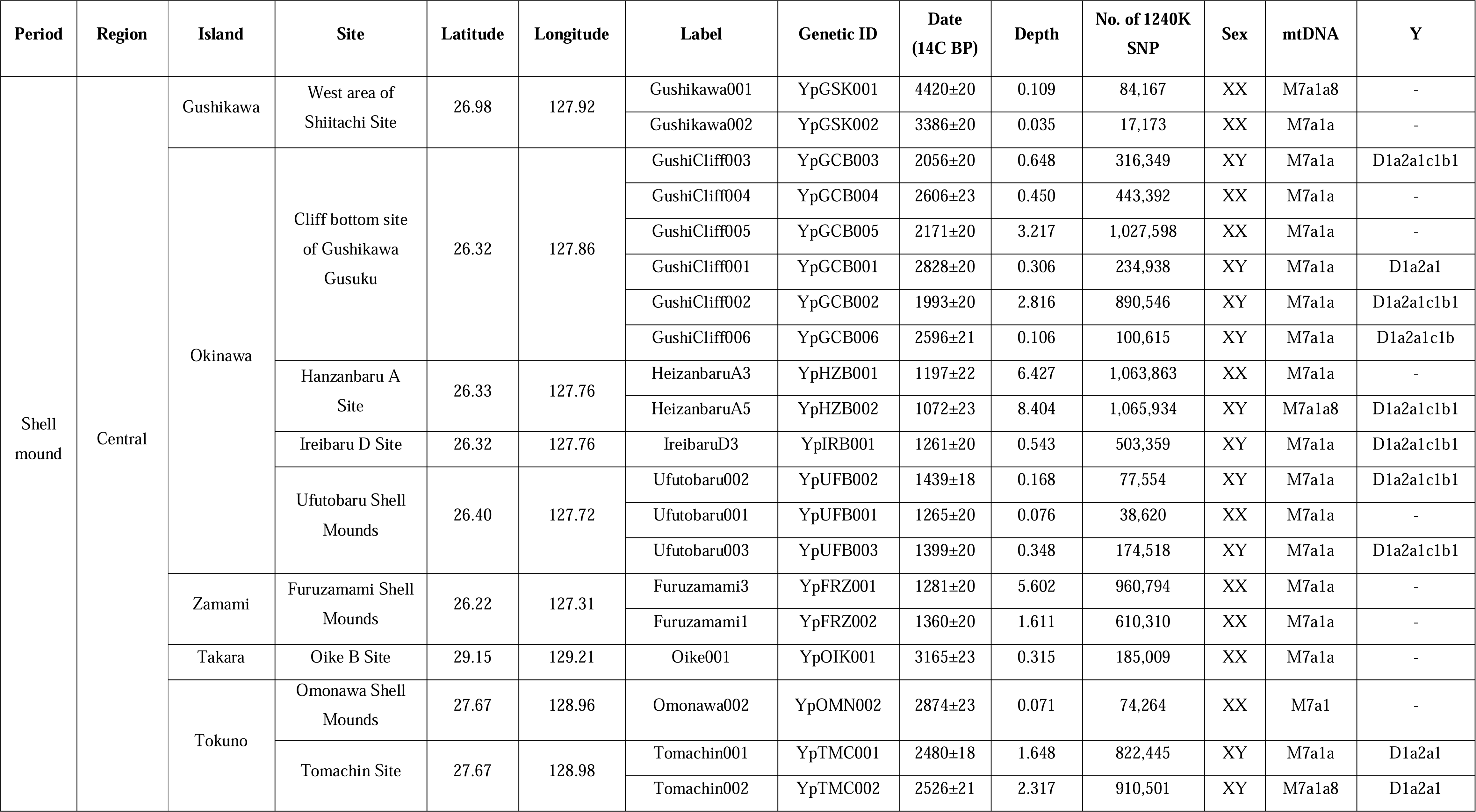
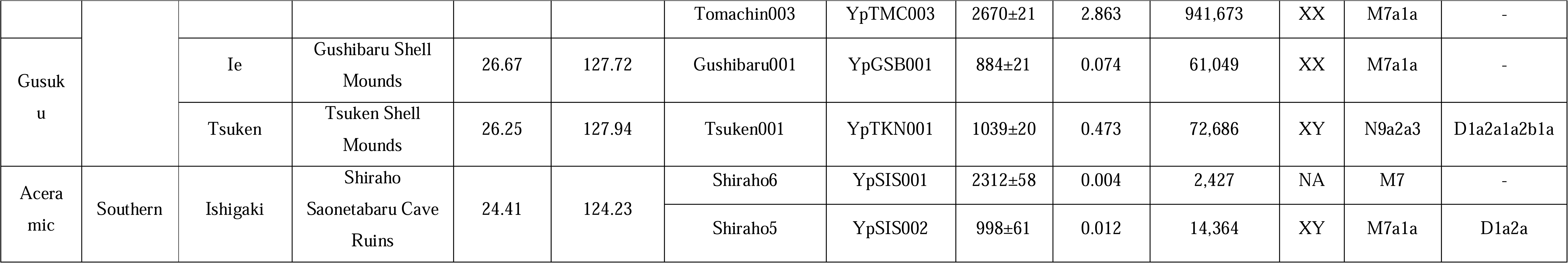
Detailed information for newly sequenced Ryukyu ancient genome.

Except for Shiraho6 from the Shiraho Saonetabaru Cave site, the sequence quality of the samples was sufficient for population genomics analyses (Supplementary Fig. 3): depth > 0.01 with more than 10K SNPs. Most samples showed a low contamination rate (less than 0.09%), but due to the low sequencing depth, the contamination rate could not be estimated for Shiraho6 and Shiraho5 (Supplementary Table 2).

We investigated kinship among the Ryukyu Jomon individuals using identity by descent (IBD) segment sharing (Ringbauer et al. 2024) and found evidence of close genetic relationships among the three individuals from the Tomachin site on Tokuno-shima Island from ∼2,500 YBP (Supplementary Fig. 4). Based on the total length of IBD segments and the count of IBD segments >8cM, we estimated that Tomachin003 and Tomachin001 were siblings, while Tomachin002 was the son of Tomachin001. Consequently, Tomachin003 was thought to be the aunt of Tomachin002.

We also found multiple pairs of individuals who were not first- or second-degree relatives but who shared more than 2 segments with length of >20cM in length, potentially indicating a more distant kinship (third-degree plus). These distant kinship relationships were inferred within the Gushikawa site and the Heizanbaru site on Okinawa Island. Interestingly, we also inferred such links between different sites. HeizanbaruA3 (1,200 YBP) has distant kinship with Ufutobaru003 (1,400 YBP) from a separate site on Okinawa Island and with Furuzamami3 (1,200 YBP) on Zamami Island. This indicates a certain degree of local migration among the Ryukyu Jomon, including between islands.

In analysis of runs of homozygosity (ROH) segments using hapROH (Ringbauer et al. 2021), the length of the ROH varied significantly among the ancient individuals, from 4 to 286 cM. ROH segments longer than 12cM are indicative of recent inbreeding, as observed in the Tomachin individuals and the IreibaruD3 individual on Okinawa Island.

We determined mitochondrial DNA and Y chromosome haplogroups from genotyping data. All of the Ryukyu Jomon individuals had a M7a haplogroup for mitochondrial DNA and all male individuals showed a D1a2a1 haplogroup for Y chromosome (Table 1).

### Structures of ancient and modern Ryukyu populations

We obtained WGS data for 273 modern individuals collected across the Ryukyu Archipelago: Okinawa Island and its surrounding islands (OKI, n = 97), Kume Island/Tonaki Island/Awa Island (KME, n = 60), the Miyako Islands (MYK, n = 50), the Yaeyama Islands (YAE, n = 48), and the Amami Islands (AMA, n = 18).

Obtained Ryukyu WGS data was combined with other genomes to infer the population structure using a principal component analysis (PCA). We extracted 1240K SNPs from newly sequenced and previously reported ancient genomes (Supplementary Tables 3 and 4) and projected these data into a PCA generated by modern genome data. When we focused on East Asian populations (Fig. 1C), three clusters (Hondo, Ryukyu, and Jomon) were aligned along the PC1 axis. Within the Jomon cluster, Hondo and Ryukyu Jomon exhibited slightly different PC1 scores, suggesting genetic differentiations between Hondo and Ryukyu Jomon. Modern individuals from different island groups in the Ryukyu Archipelago formed a cluster as shown in Fig. 1C. When we plotted another PCA result using only Japanese populations (Fig. 1D), Ryukyu and Hondo Jomon individuals were more clearly differentiated on the PC1 axis, and the PC2 axis was mainly explained by the genetic diversity within the Miyako Islands, as previously reported (Matsunami et al. 2021; Koganebuchi et al. 2023). For the Ryukyu Jomon population, linear regression analyses indicated that PC1 values are associated not with location of the site (Supplementary Fig. 5A; R^2^ = -0.0264, p = 0.4738) but with date (Supplementary Fig. 5B; R^2^ = 0.1658, p = 0.0472). However, sequence depth was also associated with PC1 values (Supplementary Fig. 5C; R^2^ = 0.2388, p =0.0195). Using imputed ancient data, we also observed that Ryukyu and Hondo Jomon clusters were aligned along the PC1 axis (Supplementary Fig. 6).

ADMIXTURE analysis indicated that three ancestral populations (K = 3) best explain the genetic profiles of East Asian populations. These ancestries seem to correspond to Jomon, northern continental, and southern continental lineages (Fig. 1E; Supplementary Fig. 7). The Hondo and Ryukyu Jomon shared a single ancestry, and differentiation between the Jomon populations was not observed in the results of ADMIXTURE.

### *F*-statistics between Ryukyu and other populations

Pairwise outgroup f3 statistics revealed that the Ryukyu Jomon have relatively high genetic affinities with surrounding modern and ancient East Asian populations (Supplementary Fig. 8; Supplementary Tables 5 and 6). The dendrogram for modern and ancient Japanese individuals based on individual-level pairwise f3 statistics shows two distinct clusters (Fig. 2A): one consisting of Jomon individuals and the other of non-Jomon individuals. The Jomon cluster was subdivided into two groups, one mainly corresponding to the Ryukyu Jomon individuals and one mainly corresponding to Hondo Jomon individuals. These results were replicated using population-level f3 statistics (Supplementary Fig. 9). We also found that ancient individuals from the Aceramic culture in the Southern Ryukyus were included in either the Hondo Jomon cluster or the non-Jomon cluster (Fig. 2A). However, since the data quality for the Aceramic individuals was very low (Supplementary Fig. 3), the results need to be carefully interpreted.

**Fig. 2:**
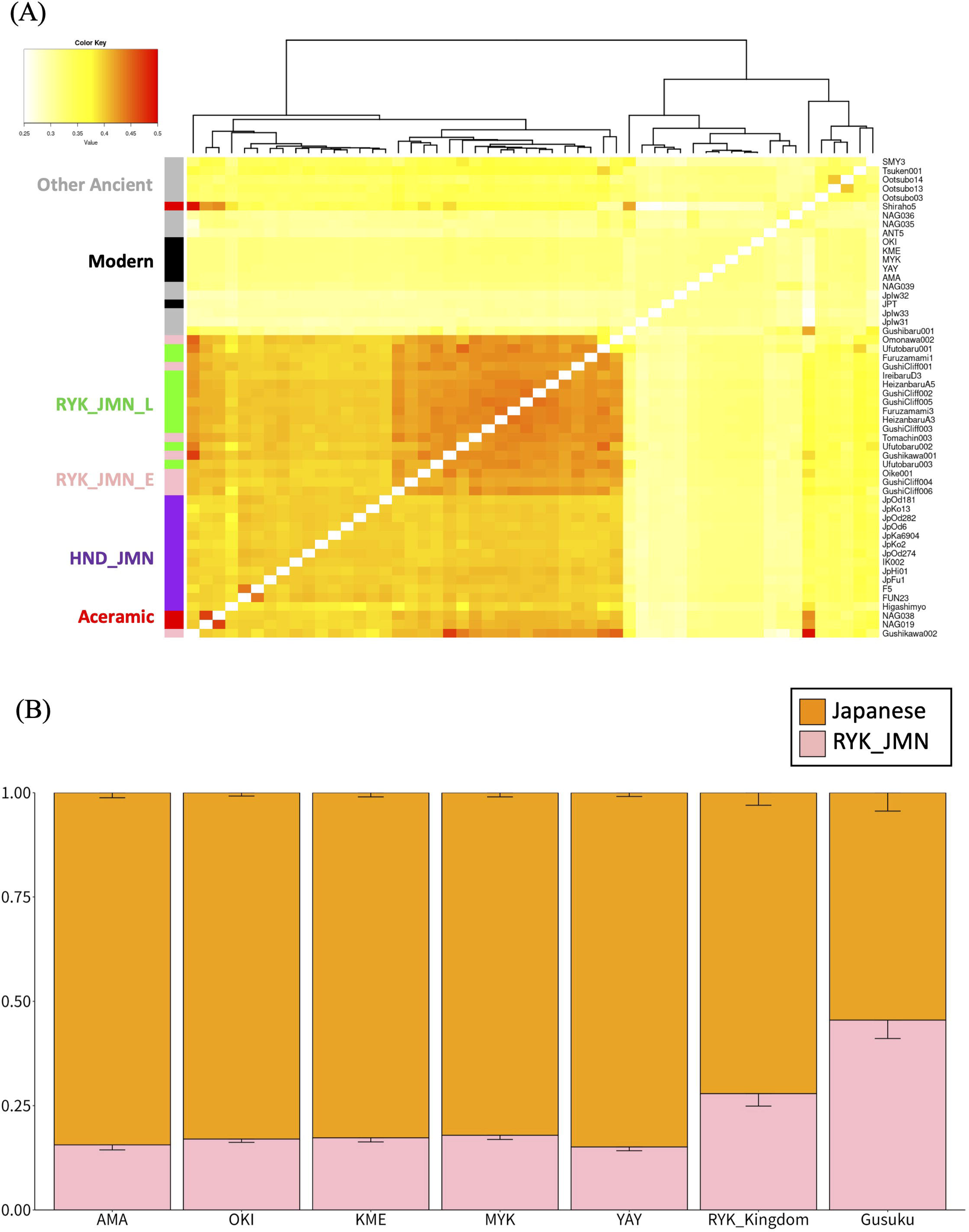
Genetic affinity between ancient/modern Ryukyu and other populations. **(A)** Heatmap of outgroup f3 statistics between Japanese modern populations and ancient individuals. High f3 statistics indicate high genetic affinity. **(B)** Ancestry estimation using qpAdm. Target populations are modern and ancient Ryukyu, and Japanese and Ryukyu Jomon were used for source populations. HND_JMN: Hondo Jomon, RYK_JMN_E: Early Ryukyu Jomon, RYK_JMN_L: Late Ryukyu Jomon.

Next, we calculated the f4 statistics to assess relative Jomon ancestry in modern populations: f4(Mbuti, Jomon; OKI, JPT). The results indicated that the genetic contribution from the Jomon population is greater in the modern Ryukyu population (OKI) than in the modern Hondo population (JPT) (Supplementary Fig. 10). Although both Hondo and Ryukyu Jomon individuals showed a genetic affinity with OKI, Ryukyu Jomon individuals tended to show greater absolute f4 statistics than Hondo Jomon individuals (Supplementary Fig. 10). We did not detect any evidence of variation in gene flows from the Jomon population into modern Ryukyu populations (OKI, MYK, YAE, AMA, and KME) (Supplementary Table 7).

We also calculated f4(Mbuti, X; Hondo Jomon, Ryukyu Jomon) to detect the relative gene flow from other ancient and modern Asian populations (X) into two Jomon populations. As expected, highly positive values of the f4 statistics were observed for modern Ryukyu populations assigned to X. This indicated that modern Ryukyu populations inherited relatively high proportion of genetic components from the Ryukyu Jomon (Supplementary Fig. 11A). Unexpectedly, the f4 statistics also showed significantly positive values (Z > 3) when we assigned modern or ancient Asian populations outside the Japanese Archipelago to X. This suggested that compared with the Hondo Jomon, the Ryukyu Jomon received a slightly but significantly higher level of gene flow from another population (Supplementary Fig. 11B). We further evaluated the difference of such gene flow between the Early and Late Ryukyu Jomon periods (before and after 2500 YBP) using f4(Mbuti, X; Early Ryukyu Jomon, Late Ryukyu Jomon). However, no significant f4 statistics were detected for any modern or ancient Asian population tested as X (Supplementary Tables 8 and 9). From a prehistoric archaeological perspective, one of the candidates for the source of the gene flow may be Upper-Paleolithic people in the Ryukyu Archipelago. Although no data are available for Upper-Paleolithic Ryukyu people, an Upper-Paleolithic individual of China, Tianyuan, showed a non-significant f4 statistics (Supplementary Fig.11) and thus was not a plausible source.

### Local Admixture in the Ryukyu Archipelago

Using the f4 ratio test, we estimated the admixture proportion between the Jomon people (Hondo or Ryukyu Jomon) and continental migrants (CHB or Korean) during formation of the Japanese populations. The results of the f4 ratio test indicated that the admixture proportions from Jomon people to the modern Ryukyu populations were 0.159-0.279, whereas those to the modern Hondo population (JPT) were 0.035-0.136 (Supplementary Fig. 12; Supplementary Table 10). When CHB, instead of Korean, was used as the continental parental population, the estimated admixture proportion from Jomon people was increased.

Next, we applied the qpWave and qpAdm algorithm to estimate the admixture proportion in the ancient and modern Ryukyu populations. The results of qpWave calculation indicate that at least two migration waves to the Ryukyu Archipelago are required to explain the past and present population structures (Supplementary Table 11).

Since the results of the f4 statistics suggested the presence of the gene flow from a population outside the Japanese Archipelago into the Ryukyu Jomon, we attempted to identify the source population that contributed to the admixture. Using two-way qpAdm with Hondo Jomon as one of the source populations, many ancient and modern Asian populations were identified as potential source populations for the Ryukyu Jomon (Supplementary Tables 12 and 13). From a geographical perspective, Austronesian-speaking people living in Taiwan and the Philippines are candidates for the source population. While indigenous Taiwanese, Ami and Atayal, were acceptable as sources in the two-way qpAdm algorithm, the continental populations such as Oroqen, Hezhen, Ulchi, Daur, Korean, and northern Han exhibited a greater tailed p-value (Supplementary Table 13). To determine which of the Austronesian-speaking or continental populations are a more plausible source population, we further conducted a rotated qpAdm analysis using Ami and Oroqen as reference populations. The results indicate that Oroqen (admixture rate = 0.176, tail p-value = 0.54) is more likely to serve as a source population than Ami (admixture rate = 0.104, tail p-value = 0.04) (Supplementary Tables 14), which is consistent with the Ryukyu Jomon receiving gene flow from a continental, but not insular, Asian population.

We also modeled the formation of the Ryukyu populations by admixture between the Ryukyu Jomon and modern Hondo Japanese (Supplementary Table 15; Fig 2B). The results showed that the proportion of Jomon ancestry gradually decreases over time (Gusuku > Ryukyu Kingdom > modern). The admixture rate from the Ryukyu Jomon to modern Ryukyu populations ranged from 0.151 to 0.173 and remained largely stable testing each island group (OKI, MYK, YAE, AMA, and KME) as the target population, although it was slightly lower for YAE and AMA.

Phylogenetic trees were constructed using the Treemix program (Pickrell et al. 2012; Supplementary Fig. 13). We confirmed that the Ryukyu and Hondo Jomon cluster together and that all of admixture edges were directed from the ancient Jomon branches to the modern Japanese branches, as expected. Additionally, we reconstructed a model of population formation using qpGraph. Although we attempted to introduce an admixture between the Jomon and continental populations to form the Ryukyu Jomon, the proportion from the continental population was estimated to be zero (Supplementary Fig. 14). Consequently, the resulting model took the form shown in Fig. 3, although the worst z score in the f4 statistics was 4.412. The admixture rate from the Hondo Jomon to the modern Japanese was estimated to be 13% and that from the Ryukyu Jomon to the modern Ryukyu population was 18%.

**Fig. 3:**
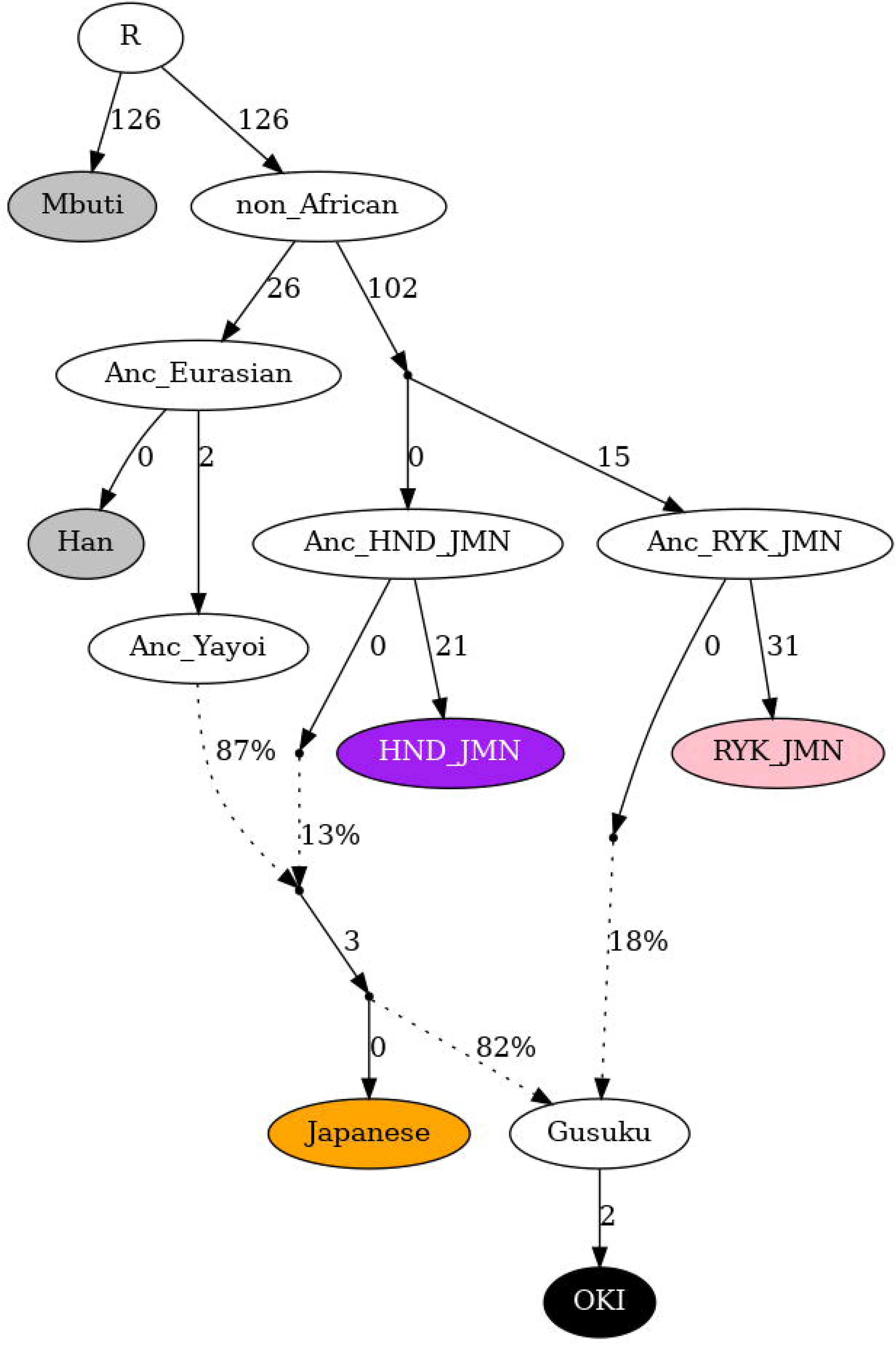
Admixture graph modeling for Ryukyu populations. Based on f-statistics analysis, a suitable evolutionary model for modern and ancient Ryukyu population was proposed (|Z| = 4.412). In this model, two ancestral Jomon populations were assumed, and these populations were split from the ancestral non-African population. Colored nodes represent actual populations as input data, whereas others are hypothetical populations.

### Demographic dynamics for the Ryukyu populations

To understand the demographic dynamics of the Ryukyu populations in detail, we conducted further population genomics analyses using imputed ancient genomes. Because we could not obtain genomic data of good enough quality from ancient samples from the Southern Ryukyus, we only used samples from the Central Ryukyus in the subsequent analysis.

We estimated the date of the admixture between Jomon and CHB as being representative of the continental population using the software ALDER (Loh et al. 2013). The date inferred by ALDER tends to indicate the most recent admixture event between the two ancestral components. Therefore, it should be noted that the admixture may not have been a single event and that the admixture event does not necessarily represent direct mixing between Jomon and CHB. The estimated date of Jomon admixture in the Hondo Japanese (JPT) population was 54.49 ± 2.37 generations ago (Fig. 4A), which corresponds to about 1,526 YBP during the Kofun period when we consider 28 years/generation. In contrast, the estimated date of Jomon admixture in the Ryukyu population (OKI) was 36.77 ± 1.71 generations ago (approximately 1,030 YBP) (Fig. 4B). The dates of Jomon admixture were similar among island groups in the Ryukyu Archipelago (Supplementary Fig. 15; Supplementary Table 16).

**Fig. 4:**
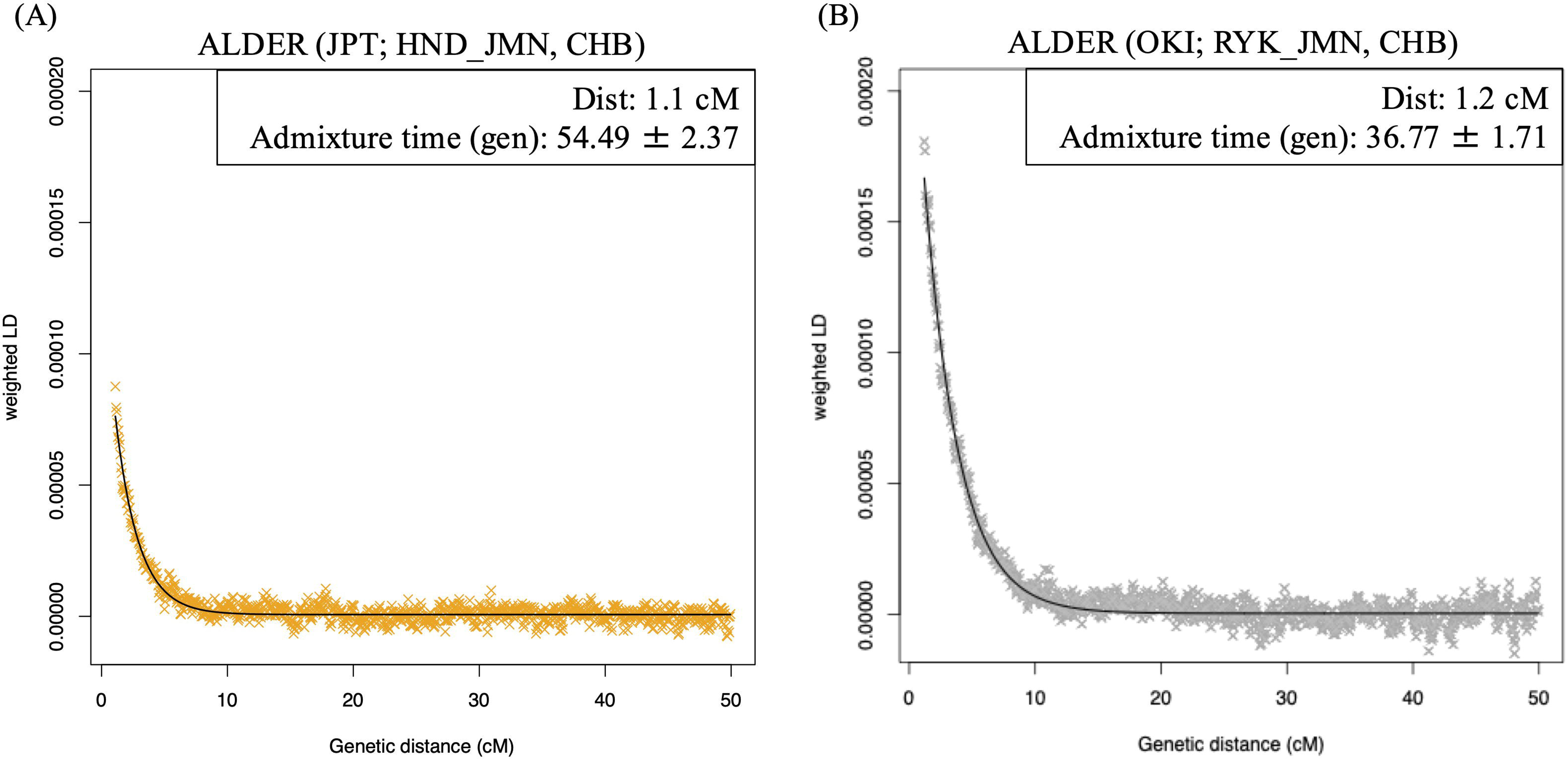
Admixture dating between JPT/OKI and Jomon ancestry. **(A)** LD decay curve for JPT using Hondo Jomon and CHB as sources. **(B)** LD decay curve for OKI using Ryukyu Jomon and CHB as sources. The fitted trendline considers a minimum distance of 1.1 and 1.2□cM for JPT and OKI populations, respectively.

As a signature of a population bottleneck or prolonged small effective population size, we estimated the number and length of ROH for each modern and ancient population using the imputed data. From the distribution of ROH ranging from 1-5 Mbp, the Hondo Jomon showed longer ROH than the modern Japanese (Fig. 5A). In addition, the Ryukyu Jomon exhibited longer ROH than the Hondo Jomon, suggesting that the Ryukyu Jomon had experienced a more severe bottleneck (t = 2.989, df = 18.859, *P* = 0.008; HND_JMN N=14, RYK_JMN N=19). In contrast to the distribution of total number and total length of ROH (Supplementary Fig. 16A), no difference was observed in the distribution of ROH ranging more than 5 Mbp (Supplementary Fig. 16B).

**Fig. 5:**
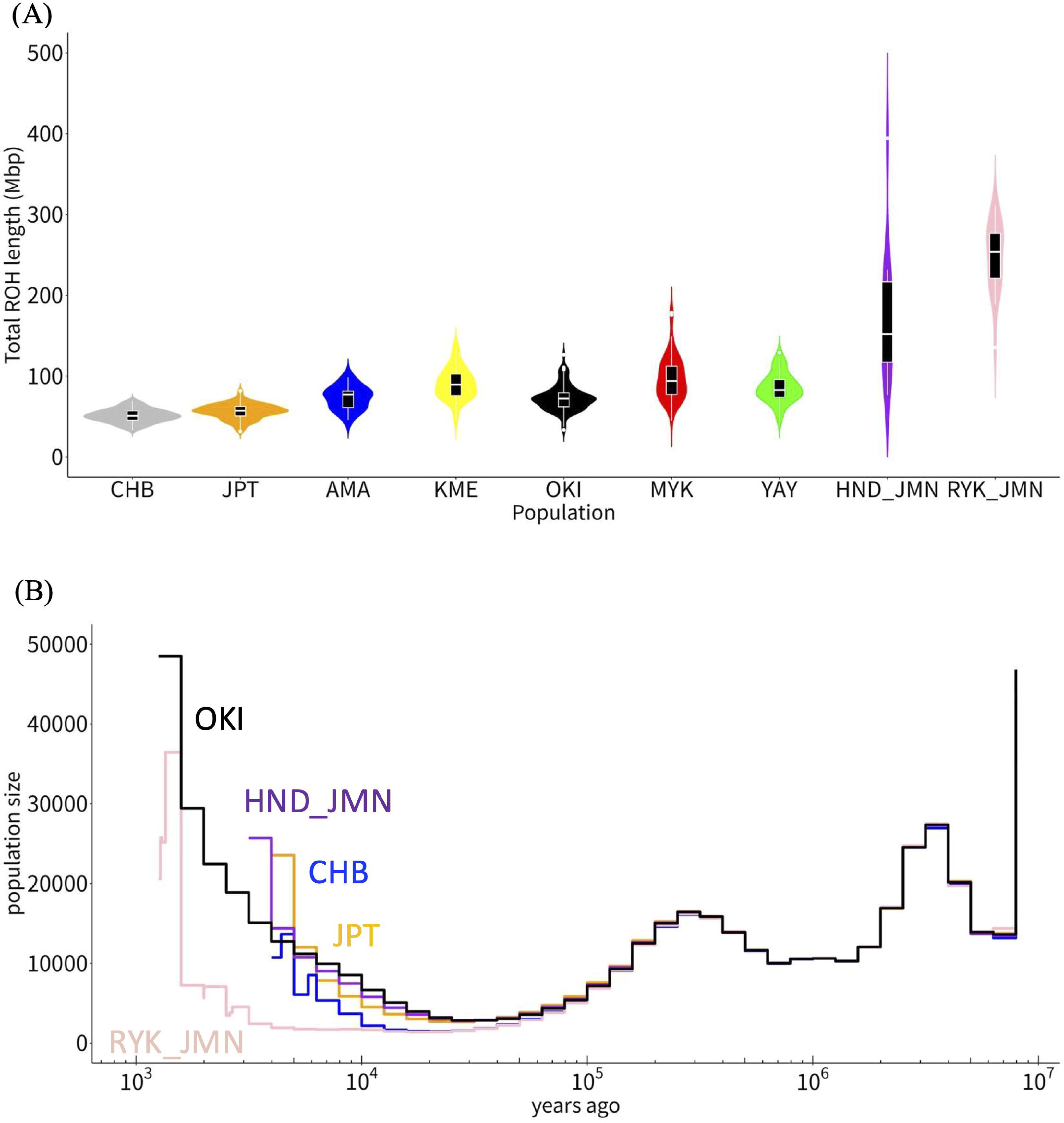
Inferences of population size using haplotype data. **(A)** Violin plot showing distribution of total length of short ROH (1-5 Mbp) for each population with rotated kernel probability density. Each black box indicates the interquartile range and each horizontal white line indicates the median. **(B)** Changes in effective population sizes were estimated using RELATE. The vertical axis represents population size with a common logarithmic scale. The horizontal axis represents generation times. We assumed 28 years per generation.

We also conducted an ancestral recombination graph analysis using RELATE to estimate the historical change in population size (Speidel et al. 2019; Fig. 5B). As with the ROH analysis, the results indicated that the Ryukyu Jomon had a severely decreased population size. The demography is mostly identical between the Ryukyu and Hondo Jomon populations to about 10,000 YBP. Thereafter, these two populations seem to have been genetically differentiated. We further conducted a time-stratified f4 ratio test using RELATE and twigstats (Speidel et al. 2025; Supplementary Fig. 17; Supplementary Table 17). However, no clear migration signal from Jomon population to other Japanese population was detected when either time threshold (500 or 300 generations) was applied.

Finally, we estimated divergence time and the population sizes of the Ryukyu and Hondo Jomon populations based on coalescent simulation using fastsimcoal2 (Excoffier et al. 2021). Considering the results of the ancestral recombination graph, we simplified the model assuming that these two populations split after the population bottleneck during the Out-of-Africa migration, and both kept a constant population size without migration to each other (Fig. 6A). The joint fold site frequency spectrum (SFS) was used as the summary statistic in the simulations. The observed SFS, the expected SFS for the best-fitting model, and the Anscombe residuals were shown in Fig. 6B. The results of our simulations suggested that the timing of the split between the Ryukyu and Hondo Jomon populations was 246 generation ago (95% CI: 230-265), corresponding to 6,900 YBP (28 years/generation), and the population size of Ryukyu Jomon was 2,099 (95% CI: 2,026-2,296). Since the model does not take admixture or migration into account, the estimated divergence time is likely to represent the lower bound.

**Fig. 6:**
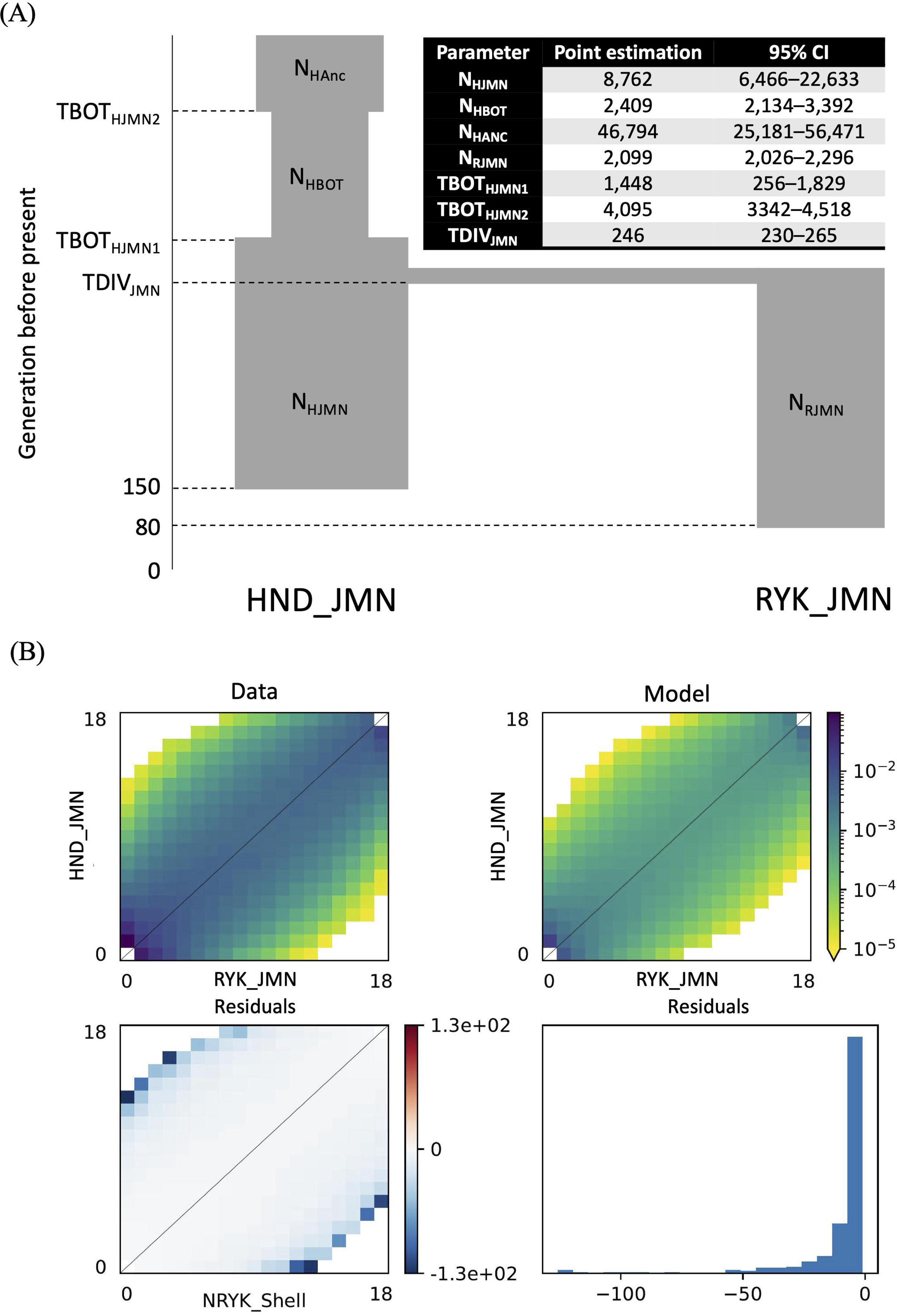
Proposed evolutionary model for coalescent simulation. A simple demographic model was proposed for coalescent simulation using fastsimcoal2, considering the results of other analysis. **(A)** We estimated all parameters shown in the figure, such as effective population sizes and divergence times. The resulting point estimations with 95% confidence interval of coalescent simulation are shown in table. **(B)** Observed and expected joint fold SFS between Hondo and Ryukyu Jomon populations with Anscombe residuals between observed and expected SFS were plotted.

### Jomon-derived regions within the modern Japanese genome

We calculated F_ST_ for each locus between the Ryukyu and Hondo Jomon populations and found that 19 loci showed F_ST_ > 0.7. These loci are included in functional genes, such as SSBP3, FAM180A, and AXCR1C (Supplementary Table18).

To understand the details of the Jomon components in modern populations, we decomposed admixed chromosomes using a local ancestry inference. We estimated haplotypes derived from the Jomon population in both modern Ryukyu and Hondo individuals and calculated the rate of Jomon ancestry for each chromosomal region in each population (Fig. 7A). The Jomon rates in each chromosomal region were clearly correlated between JPT and OKI (Fig. 7B; r = 0.568, t = 41.363, df = 4600, P < 2.2 ×10^−16^) and there was no region with a signature indicating different selection pressures between the populations. Our analysis suggested that rates of Jomon ancestry in modern Ryukyu genomes were relatively higher than those in modern Hondo genomes on average (mean of difference = 0.106). In particular, we found that a locus showing a high Jomon ancestry rate is located near the telomeric region of chromosome 14 (Supplementary Tables 19 and 20; chr14:105,562,800-106,883,658). This region includes the gene cluster encoding the immunoglobulin heavy chain.

**Fig. 7:**
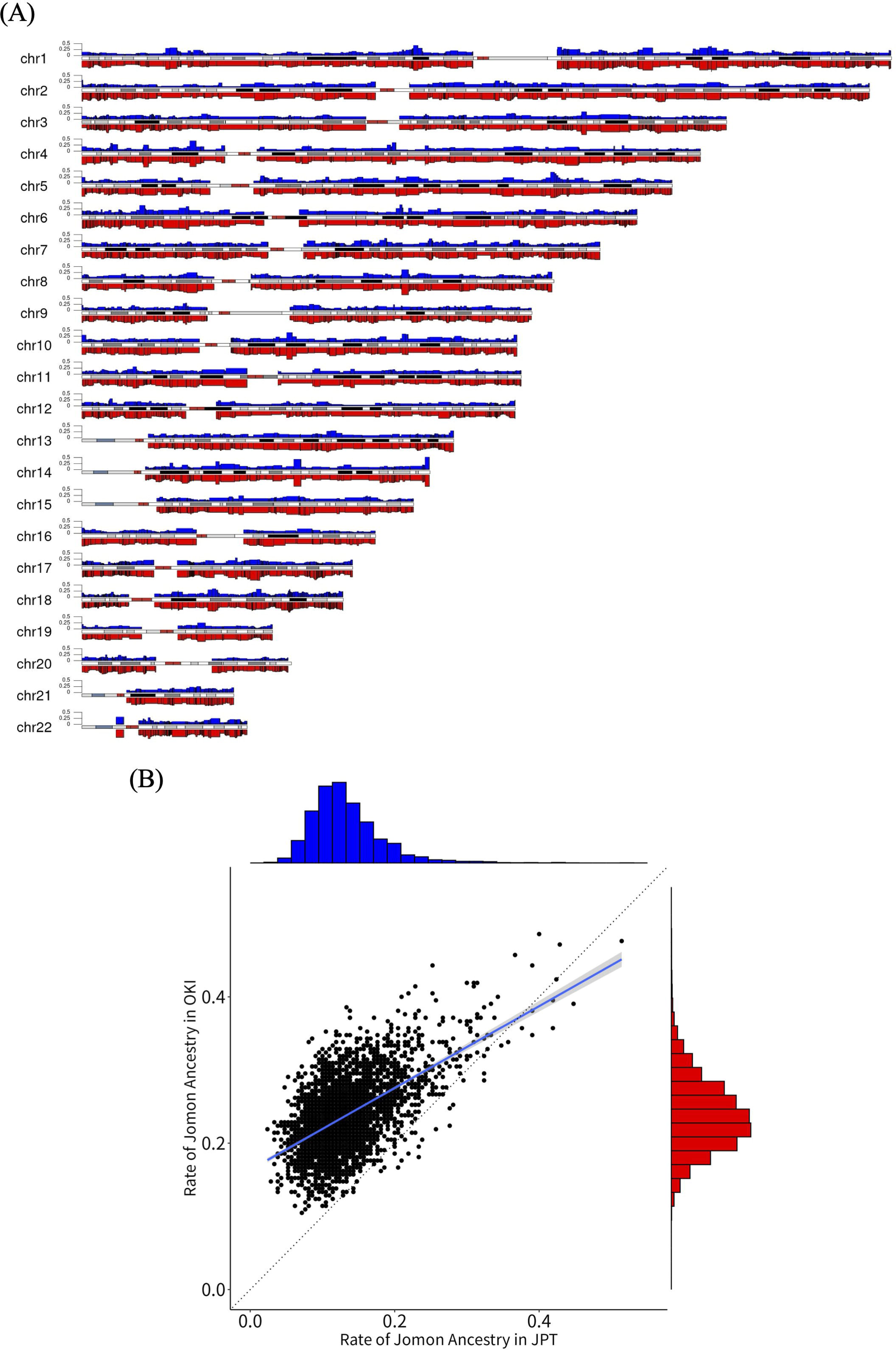
Signatures of Jomon-derived haplotypes in modern Hondo and Ryukyu genomes. **(A)** Chromosomal overview of the distribution for the rate of Jomon ancestry in the JPT/OKI populations. The upper track shown in blue is the rate of Jomon ancestry for each haplotype in the JPT population, while the lower track shown in red is the rate of Jomon ancestry for each haplotype in the OKI population. (B) Scatter plot of rate of Jomon ancestry between JPT and OKI populations. The regression lines are shown with standard errors. We observed a significant correlation between the rate of Jomon population from JPT/OKI (r = 0.568, t = 41.363, df = 4600, P < 2.2 ×10^−16^).

## Discussion

In the dual structure model, Hanihara (1991) argued that the Japanese Archipelago was occupied by the indigenous Jomon people and that differences between the Hondo and the Ainu/Ryukyu populations were caused by migrations from the East Asian continent, predominantly into the Hondo region. Technological advances make it possible to test these hypotheses using genomic data. Genome-wide SNP studies demonstrated that, although there is some genetic similarity between the Ainu and Ryukyu populations, the Ryukyu population are genetically closer to the Hondo Japanese than to the Ainu (Jinam et al. 2012). Building on the basic concept of the dual structure model, researchers recently hypothesized that assuming additional migrations is more suitable for explaining the genomes of the ancient and modern Japanese (Cooke et al. 2021; Jinam et al. 2021; Liu et al. 2024). This study focusing on the Ryukyu Archipelago highlights that the origins of the Japanese population may be more complicated than previously thought, and the modern Ryukyu and Hondo people do not simply share the same Jomon ancestors.

We demonstrated that the Ryukyu and Hondo Jomon populations are genetically differentiated (Figs. 1C, D and 2). A previous study has suggested that Hondo and Ryukyu Jomon individuals have a different facial morphology of the skull (Fukase et al. 2012). In this study, we also detected genetic polymorphisms that are highly differentiated between the Hondo and Ryukyu Jomon populations (Supplementary Table 18), which can be candidates for the loci that explain phenotypic differences between populations.

Using coalescent simulation, we estimated the divergence time between the Ryukyu and Hondo Jomon to be 6,900 YBP. Interestingly, this date coincides with the emergence of Tsumegata-mon (Fingernail-impressed) pottery culture associated with the onset of full-scale stone tool production in the Central Ryukyus (Takamiya et al. 2019; Pearson 2013). This culture emerged after the Kikai-Akahoya eruption (7,300 YBP) (Uchiyama et al. 2023) and is considered to have been independent of the Tsumegata-mon culture in the Hondo region that emerged during the Incipient Jomon period (before 11,500 YBP). Thus, the origin of the Tsumegata-mon pottery culture in the Central Ryukyus is still unclear. Although Jomon pottery in the Central Ryukyus was discovered sporadically from earlier periods (Yamasaki 2020), archaeologists have regarded the Tsumegata-mon pottery culture as the establishment of the Ryukyu Jomon period, based on increased evidence of human settlement. Although our simplified simulation model, which assumes no migration or admixture, may under- or over-estimate the divergence time, the estimation does not contradict our understanding that people migrated from Hondo to the Central Ryukyus and introduced Jomon pottery cultures.

Previous morphological studies of ancient bones have suggested that the Upper-Paleolithic Ryukyu people were distinct from the Jomon people and were likely derived from prehistoric Southeast Asian populations with Australo-Melanesian affinities (Kaifu et al. 2011). Previous studies have suggested that the mtDNA of the Upper-Paleolithic Ryukyu individuals does not show continuity with that of subsequent Jomon individuals or present-day populations in the same region (Mizuno et al. 2021; Shinoda et al. 2013b). Consistent with those findings, this study found no evidence that continental Upper-Paleolithic people represented by Tianyuan contributed genetically to the Ryukyu Jomon. Instead, our f4 and qpAdm analyses detected that the Ryukyu Jomon had received a slightly greater genetic contribution from another population than the Hondo Jomon had, demonstrating that the source population is likely a northern continental group but not the indigenous Taiwanese. At this moment, we cannot conclude whether gene flow occurred directly from the Asian continent or through Hondo. Although previous studies have demonstrated that the genomes of historical and modern Ryukyu individuals harbor continental genetic components (Cooke et al. 2023; Liu et al. 2025), the introduction of certain components into the Ryukyus may date back to an earlier era than the Late Ryukyu Jomon (Yayoi) period.

Our findings suggest that the Ryukyu Jomon experienced a severe genetic bottleneck after the divergence from the Hondo Jomon and maintained a small population size (about 1,700 in RELATE and 2,100 in fastsimcoal2) for more than 5,000 years. Recently, Kawai et al. (2023) estimated the transition in population size from ancient times to the present using whole genome data for modern Ryukyu individuals. However, the previous estimation may not directly reflect the population size of the Ryukyu Jomon because the large proportion of the modern Ryukyu ancestry is derived from the historic Hondo population. Therefore, the present study provides new evidence on population size changes in the Ryukyu Archipelago during the prehistoric period using genomic data from Ryukyu Jomon individuals.

The origins of prehistoric cultures in the Southern Ryukyus, the Shimotabaru and Aceramic cultures, remain poorly understood (Yamagiwa et al. 2019). Some archaeologists consider that these cultures originated in Taiwan or the Philippines. However, previous genome analysis demonstrated that ancient individuals from the Aceramic period on Miyako Island have a genetic background similar to that of the Jomon (Robbeets et al. 2021). In the present study, we attempted to add additional ancient samples from the Southern Ryukyus; however, high-quality genomic data could not be obtained. In the analysis using f3 statistics (Fig. 2), individuals from the Southern Ryukyus in the Aceramic period clustered together with the Hondo Jomon individuals. Nonetheless, due to the limited quantity and quality of the data, it would be premature to conclude that the aceramic culture was introduced from the Hondo region, skipping over the Central Ryukyus. Therefore, collection of additional high-quality data and comprehensive data analyses are required to clarify the origins of the prehistoric cultures in the Southern Ryukyus.

From an archaeological perspective, cereal farming and the use of iron tools were introduced to the Central Ryukyus between the 8th and 12th centuries, after which the Ryukyu Archipelago was considered to have entered the Gusuku period (Takamiya and Nakamura 2021). Clarifying whether this cultural transition resulted from the transmission of cultural practices alone or from human migration is also archaeologically important. As previously reported (Koganebuchi et al. 2023), the present study showed that the genetic contribution of migrants from the Hondo region is substantial in the modern Ryukyu population (83% and 82% in OKI by qpAdm and admixture graph, respectively). Although the admixture proportions were largely stable across different island groups, it is noteworthy that the Yaeyama Islands, despite being the farthest west, showed a relatively high proportion of Hondo-derived admixture. Furthermore, we showed that ancient individuals from the Gusuku period and the Ryukyu Kingdom period have lower proportions of Hondo-derived admixture. This suggests that migration and admixture events did not occur at a single point in time, but rather accumulated gradually over time.

We aimed to estimate the timing of the admixture between the indigenous Ryukyu Jomon and the Hondo-derived immigrants after the Gusuku period. ALDER likely failed to make this determination because the modern Hondo population containing some Jomon ancestry is genetically too close to the modern Ryukyu populations (Loh et al. 2013). Therefore, we instead used CHB as one of source populations and detected the admixture between the Jomon and continental genomic components. The Jomon admixture in the modern Ryukyu populations was estimated to be approximately 1,000 YBP (Fig. 3), which corresponds to the timing of the introduction of agriculture. This is also consistent with a previous estimation using individuals from the Ryukyu Kingdom period on the Miyako Island (historical Nagabaka) (Cooke et al. 2023). These results suggest that the introduction of agriculture to the Ryukyu Archipelago was accompanied by human migration.

We showed that the rates of Jomon ancestry in each chromosomal region were higher in OKI than in JPT and were clearly correlated between the modern populations (Fig. 7). Concurrently, we did not observe any genomic regions with distinctly higher or lower levels of Jomon ancestry that were specific to OKI. The genomic regions showing higher levels of Jomon ancestry in both populations may be candidates for adaptive introgression, but a comprehensive understanding of this phenomenon will require elucidating the associations between phenotypes and genetic polymorphisms, as well as the functions of nearby genes.

Although our study provides new insights into the demography of Japanese populations, we should also recognize its limitations. First, due to the small sample size and the low quality of the ancient genome samples, we could not conduct a detailed analysis of the Southern Ryukyus. Second, because of the absence of genome data from Paleolithic individuals, we could not explicitly clarify the relationships between the Paleolithic people and the Ryukyu Jomon. Third, although archeological evidence indicates that trade between the Ryukyus and external region increased during the Late Ryukyu Jomon period in the Central Ryukyus (Kinoshita 2012), no genetic traces of this were detected. Fourth, we could not decompose admixture events during and after the Gusuku period. Additional genome information from the Ryukyus will facilitate the resolution of these remaining questions.

## Materials and Methods

### Ethics statements

Prior to sharing modern genome information used in this study, all participants provided written informed consent. The study protocol was approved by the Ethics Committee of the National Center for Global Health and Medicine (Approval No. NCGM-S-003460-06) and the Ethics Committee on Life Science and Medical Research Involving Human Subjects, University of the Ryukyus: Approval No. 18-2300-07-00-00, No. 21-1826-17-03-00 (for modern genome data) and No. 23-2213-00-00-00 (for ancient genome data).

### Modern sample collection and sequencing

A total of 273 individuals living in the Ryukyu Archipelago (OKI: n = 97, KME: n = 60, MYK: n = 50, YAE: n = 48, and AMA: n = 18) participated in this study and gave written informed consent before their enrollment. The origin of individual participants was determined using information on the birthplace of their four grandparents obtained by questionnaire. Genomic DNA was extracted from saliva or blood samples. WGS was conducted at Macrogen Japan Corp. (Tokyo, Japan) and Genesis Healthcare Co. (Tokyo, Japan). DNA quality was assessed using the picogreen method, and the condition of the DNA was assessed by gel electrophoresis. A WGS library was constructed using a TruSeq DNA PCR-Free Library Preparation kit (Illumina Inc., San Diego, CA, USA) according to manufacturer protocols. We sequenced the DNA using 2□×□150-bp paired-end reads on a HiSeq X sequencing platform (Illumina Inc.).

### Ancient sample collection and genomic mapping

Excavated ancient bones (25 samples from 12 sites) in the Ryukyu Archipelago were utilized for this study (Figure 1). Archeological evidence suggests that all of these sites date to the Shell-mound period (i.e., Ryukyu Jomon). Description of each site are provided in Supplementary Note 1. C14 radiocarbon dating was performed on the bone samples as described in Supplementary Note 2. After ancient DNA extraction and library preparation (Supplementary Note 3), DNA libraries were sequenced using HiSeq X (Illumina) with 150 base paired-end reads. The DNA sequence data were mapped to the GRCh38 reference genome using the BWA software with the mem option (Li and Durbin 2009). After removing soft clips, base recalibration was performed with the GATK4 BaseRecalibrator (Van der Auwera and O’Connor 2020). The generated bam file for each sample was utilized in further analysis.

### Molecular sex determination and contamination estimation for ancient dataset

To clarify the authenticity of the mapped sequence reads, the frequency of postmortem misincorporation and depurination, which are characteristic of ancient DNA, was checked with mapDamage2.0 (Jónsson et al. 2013). After confirming the authenticity, the contamination frequency was estimated for nuclear DNA sequences. We used ANGSD software (“MoM” estimate from “Method 1” and the “new_llh” likelihood computation) (Korneliussen et al. 2014), which estimates contamination based on the X chromosome polymorphism rate in males. Then, the molecular sex of ancient samples was determined using the method of Skoglund et al. (2013).

### Genotype calling and merge to previously reported modern and ancient datasets

We collected publicly available modern genome data from following datasets: 1000 Genomes Project Phase 3 (The 1000 Genomes Project Consortium 2012), Simon Genome Diversity Project (Mallick et al. 2016), Human Genome Diversity Project (Bergström et al. 2020), and Korean Personal Genomics Project (http://camda2021.bioinf.jku.at/kpg_prepro). We also newly obtained WGS from 273 individuals in the Ryukyu Archipelago and one previously published high-coverage Jomon sequence (F23; Kanzawa-Kiriyama et al. 2019). Raw reads of these data were mapped to the GRCh38 reference genome and variants were jointly called.

The pipeline of joint calling described in Kawai et al. (2023) was utilized for this study. Using NVIDIA Clara Parabricks (v4.1), sequence reads in fastq format were mapped to the GRCh38 reference sequence using the bwa-mem algorithm, and duplicate reads were marked. SNV and INDEL variant calls were performed using the GATK Haplotypecaller algorithm, and the output saved in genomic Variant Call Format (gVCF) files was used to generate a VCF containing genotypes from multiple samples via joint calling with scores for each variant calculated by the VQSR algorithm using the Sentienon software. The resultant variants with a genotype quality of less than 20 or fewer than 10 reads supporting the variant were subsequently filtered out. Additionally, we applied hard filtering to heterozygous genotypes where the proportion of non-reference alleles deviated significantly (p < 0.001) from 50%. After that, we removed variants that deviated significantly (p < 10^−30^) from Hardy-Weinberg equilibrium in any population or had a missing rate of 10% or higher, and only the remaining biallelic SNPs were used for subsequent analysis.

The 1240K SNPs in the Allen Ancient DNA Resource (Mallick et al. 2024) were extracted from these joint calling data. Using downloaded 1240K v54.1 dataset, positions were converted from GRCh37 to GRCh38 using the LiftOver tool (https://genome.ucsc.edu/cgi-bin/hgLiftOver). After obtaining these SNPs, quality controls of joint calling data were conducted using PLINK1.9 (Chang et al. 2015). The criteria for SNPs filtering were as follows: (1) SNP calling rates of 95% or higher, (2) shared identity-by-descent ((’) of higher than 0.5, and (3) hetero F coefficients were less than -0.07. As a result, 3,912 individuals with 1,067,494 variants were retained in the dataset. These variants were listed using mpileup in samtools with -q30 -Q20 parameters (Li 2011) and used for random haploid calling of ancient genomes using pileupCaller (https://github.com/stschiff/sequenceTools). We used samples including newly sequenced Ryukyu ancient genomes, as well as previously reported Asian and Japanese ancient genomes for calling (Table 1; Supplementary Tables 2 and 3). After calling, these modern and ancient datasets were merged.

### Mitochondrial DNA and Y chromosome haplogroups

Haplogroups of mitochondrial DNA and Y chromosome were determined. We extracted variants located in mitochondrial DNA from mpileup calling data. Then, the haplogrep program (v2.4.0) was used for the estimation of mtDNA haplogroup (Weissensteiner et al. 2016). Estimation of Y chromosome haplogroups was performed using Yleaf software (Ralf et al. 2018) with and without the -aDNA option (ignoring all G>A and C>T mutations). For further classification into Y chromosome sub-haplogroups, SNP sites in each haplogroup were called by mpileup and groups were subdivided by visually checking for the presence or absence of mutations.

### Population structure analysis

Merged data were used to examine the genetic relationships between individuals through PCA. These analyses were performed using the smartpca program in the EIGENSOFT package (Patterson et al. 2006). When ancient samples were included in the PCA, ancient samples we specified were projected to modern samples using the “poplistname” and “lsqproject” options. Ancient samples having a genomic depth of more than 0.01 were used for analysis.

To estimate individual ancestry, we conducted an ADMIXTURE analysis (Alexander et al. 2009). To decrease sampling bias, ten individuals were randomly selected from each modern population where the population includes more than ten individuals. For ancient samples, we used samples showing a depth of more than 0.01. In total, 289 individuals from the East Asian dataset were subjected to maximum-likelihood clustering analysis using an ADMIXTURE run from K = 1 to 8 with calculation of cross-validation errors for each value of K.

### *F*-statistics to detect gene flow

Individual- and population-level f3 and f4 statistics were calculated to infer gene flow between Ryukyu and other populations. For calculation of f3 statistics, we chose Mbuti as the outgroup, and pairwise comparisons between each modern and ancient Asian and Japanese populations were conducted. For calculation of f4 statistics, we also chose Mbuti as the outgroup and ancient Japanese as source populations of admixture. Then, modern Japanese were used as the target population, and the relative degrees of gene flows were estimated. The genetic differentiation among Jomon populations was estimated using f4-statistics with both Ryukyu and Hondo Jomon populations chosen as targets and other Asian ancient and modern populations selected as source populations. Both tests were carried out using admixr (Petr et al. 2019) and AdmixTools (Patterson et al. 2012).

To estimate the proportion of ancient ancestry in each modern Ryukyu population, we applied the f4 ratio test implemented in AdmixTools (Patterson et al. 2012). We selected Mbuti and CDX as the outgroup and A population, respectively. Each modern and ancient Japanese population was assigned as X population (modern: JPT, AMA, KME, MYK, OKI, and YAY; ancient: Yayoi, Kofun, Gusuku, Aceramic, and Ryukyu Kingdom). We chose modern East Asian (Korean and CHB) as the B population, and each Jomon (Hondo and Ryukyu) as the C population.

### Estimation of admixture waves

Admixture waves and genetic contributions of each ancestral population were estimated using qpWave and qpAdm in AdmixTools (Patterson et al. 2012). First, we conducted a qpWave analysis to infer the number of migration waves. We set Sardinian, Kusunda, Papuan, Dai, Ami, Naxi, and Mbuti as outgroups. We included combinations of ancient and modern Hondo populations (Japanese in SGDP/HGDP, Hondo Jomon, Yayoi, and Kofun) into the right population. Ancient Ryukyu populations (Ryukyu Jomon, and Gusuku) with or without a subset of modern and ancient Ryukyu populations (Ryukyu Kingdom, OKI, AMA, KME, MYK, and YAY) were included into the left population. Second, we conducted qpAdm analysis to estimate the proportion of genetic contributions from each ancestral population. Based on the qpWave analysis, we assumed two or three migration waves.

In addition, we conducted a two-way qpAdm analysis focusing on Ryukyu Jomon to infer the genetic contribution from populations outside the Japanese Archipelago to the Ryukyu Jomon. We set Hondo Jomon as one of the reference populations, and each Asian ancient and modern population was assigned as another reference population. Then, the genomic contributions from northern and southern populations into the Ryukyu Jomon were inferred using rotating qpAdm. At this time, we assumed Ami and Oroqen were representatives for the northern and southern populations, respectively. We calculated the admixture proportion using Hondo Jomon and these populations as references.

### Demographic inference

Bifurcation phylogenetic trees were reconstructed using Treemix (Pickrell et al. 2012). The VCF file was converted to TreeMix format using plink1.9 (Chang et al. 2015) and a python script (plink2treemix.py). We fitted 0–5 admixture edges using the Japanese ancient and modern populations and CHB was set as an outgroup.

The admixture graph was modeled using qpGraph in AdmixTools (Patterson et al. 2012). We specified parameters as outpop: Mbuti, useallsnps: YES, blgsize: 0.05, forcezmode: YES, diag: 0.0001, bigiter: 6, hires: YES, inbreed: YES. Considering previous results (Koganebuchi et al. 2023), we proposed an appropriate evolution model.

### Imputation of ancient genome

Genotyping imputation of the ancient genome was performed using GLIMPSE2 (Rubinacci et al. 2023). Before imputation, the genotypes of the modern human samples created in this study were phased using Beagle 5.4 (Browning et al. 2021). According to GLIMPSE2 instructions, the data were divided into chunks, an imputation panel was created, and the imputation was performed with bam files obtained from the ancient human analysis as input and results in VCF format. We retained ancient samples showing a depth of more than 0.1 after imputation. From these data, we filtered out SNPs showing MAF < 0.01, and individuals showing shared identity-by-descent (π̂) was higher than 0.5 for analysis of PCA, ROH, and admixture dating. After filtering, we retained 3,958 individuals with 10,880,721 SNPs.

### Inferring genomic sharing between individuals

AncIBD (Ringbauer et al 2024) was run on the imputed ancient genomes with coverage higher than 0.25 to infer IBD segments. We filtered the imputed genomes to the 1240K SNP sites lifted over to GRCh38 (Mallick et al. 2024). Allele frequency files of the imputation reference panel were created and used to make the required h5 files. AncIBD was then run in mode --IBD2 to detect both IBD1 and IBD2 segments. Only IBD calls exceeding 8cM were retained.

To infer ROH segments, we ran hapROH on the imputed ancient genomes filtered to 1240K sites (Ringbauer et al 2021). We used the 1240K 1000 Genomes reference hdf5 files and metadata in GRCh38 available from the hapROH website as a reference.

### ROH of each population

We identified ROH segments in each population using PLINK1.9 with the default setting, which selects segments containing at least 100 SNPs, and of total length ≥ 1 Mbp. We divided the resulting ROH segments into short (1-5 Mbp) and long (more than 5 Mbp) classes.

### Admixture dating

Based on the weighted linkage disequilibrium computed by ALDER v1.03 (Loh et al. 2013), admixture dates between Jomon and East Asian ancestry were estimated. We set modern Japanese (JPT, AMA, KME, MYK, OKI, and YAY) as the target population. For one-reference admixture tests, we selected Hondo or Ryukyu Jomon as the reference populations. For two-reference admixture tests, we added CHB or Korean as another target population. The default parameter setting with “checkmap: YES” was used.

### Reconstructing ancestral recombination graph using RELATE

The joint genealogies of modern and ancient individuals were estimated using the ancestral recombination graph implemented in RELATE v1.2.1 (Speidel et al. 2019). Among the imputed data, we extracted modern and ancient genomes showing a depth of more than 0.5 for later analysis. We filtered out SNPs showing an imputation posterior probability of less than 0.8 in more than 2% individuals. LiftOvered 1KGP strict mask and ancestral fasta files (https://myersgroup.github.io/relate/input_data.html) were used for analysis. After filtering, a maximum of 10 samples for each modern population, 9 samples for Ryukyu Jomon, and 12 samples for Hondo Jomon were retained for RELATE analysis. We executed RELATE with a mutation rate (m) of 1.25×10^−8^ per generation and per site, giving an effective population size of haplotypes (N) of 20,000 and specifying sample ages. Generated trees and SNP information files were used to estimate population size using the EstimatePopulationSize mode of RelateCoalescentRate.

### Time-stratified *F*-statistics using twigstats

To improve the statistical power of *F*-statistics by focusing on coalescences in recent times, we applied the time-stratified f4 ratio test using the twigstats package in R language (Speidel et al. 2025). RELATE outputs were used as inputs for this analysis. We selected Mbuti and CDX as PO and PI populations, respectively. Modern and ancient Japanese populations were assigned as PX population (modern: JPT, AMA, KME, MYK, OKI, and YAY; ancient: Yayoi and Kofun). We chose CHB and Korean as P1 populations, and each Jomon (Hondo and Ryukyu) as P2 populations. We specified a twigstats cutoff of 500 and 300 generations.

### Coalescent simulation

We conducted coalescent simulation using joint SFS as summary statistics to estimate the demography of the Jomon population. We prepared SFS from imputed genotyping data. After filtering out low-quality variants (see Reconstructing ancestral recombination graph using RELATE), we extracted Ryukyu and Hondo Jomon from genotyping data. From these data, monoallelic SNPs were removed, and the mask regions were filtered out based on 1KGP annotation. SNPs located in the protein-coding regions and CpG islands were discarded based on annotation by the ANNOVAR program (Wang et al. 2010) and the UCSC platform (https://hgw2.cse.ucsc.edu/), respectively. Ancestral and derived alleles were determined using the ancestral genome provided by The 1000 Genomes Project Consortium (2012). To reduce the computational cost, we randomly sampled 1,719,898 SNPs using vcfrandomsample function from the vcfLib library (Garrison et al. 2022). From these data, joint-fold SFS were generated by the easySFS program (Gutenkunst et al. 2009), projecting nine individuals each.

We inferred the demographic population history, namely effective population sizes, and divergence times especially focused on the Jomon population. Coalescent simulation using the fastsimcoal2 v2.7 program was carried out (Excoffier et al. 2021). A mutation rate of 1.25×10^−8^ per generation and per site was assumed. One hundred independent fastsimcoal2 runs with broad prior search ranges for each parameter were performed. Each run comprised 50 rounds of parameter estimation via the expectation/conditional maximization algorithm with a length of 100,000 coalescent simulations each. The best-fit model was selected based on the likelihood values for each run and the confidence interval for each parameter was calculated using block-bootstrapping. After the simulation, we compared the observed and expected joint-fold SFS to evaluate fitting for the proposed demographic model by dadi-cli program (Huang et al. 2023). After confirmation of divergence between Hondo and Ryukyu Jomon, we calculated the Weir and Cockerham *F_ST_* using vcftools to identify the differentiated loci (Danecek et al. 2011).

### Local ancestral inference to detect Jomon-derived regions

The Jomon-derived genomic region in JPT and OKI was inferred using local ancestral inference implemented in RFmix v2.03 (Maples et al. 2013). Because the quality and accuracy of the results depend directly on provided haplotypes, and small sample sizes often lack reference homozygote genotype calls at known variable sites, we treated the combined Hondo and Ryukyu Jomon (n = 28) as one reference population to maximize the statistical power. Since skewing of the reference sample size violates the results of local ancestral inference, we randomly selected 28 individuals from the CHB population for use as another reference population. JPT (n=106) and OKI (n=89) were selected as target populations. These data were used for input in the RFmix program. After local ancestral inference, centromeres were filtered out based on the UCSC gene annotation (https://hgw2.cse.ucsc.edu/).

We estimated the rate of Jomon ancestry for each haplotype in JPT and OKI populations from the results of local ancestry inferences. The number of haplotypes derived from Jomon in each modern population was counted. These numbers were normalized by the number of individuals in each population. We summarized the proportion of Jomon ancestry in JPT and OKI populations and plotted the results using the R package, karyoploteR library (Gel and Serra, 2017).

## Data availability

Individual modern genomic data are available in the NBDC human database (Accession: hum0562) upon review and approval. The ancient genome data used in this study have been deposited in the NCBI BioProject under accession code PRJDB40338.

## Code availability

The source code used for this study is available on github (https://github.com/mmatsunami/RyukyuJomon).

## Acknowledgements

We thank all participants who donated DNA samples for this study. We also thank the staff of the Okinawa Bioinformation Bank Project for their assistance with sample collection. Some ancient samples were provided by the Okinawa Prefecture Archaeological Center. Computations were partially performed on the NIG supercomputer at ROIS National Institute of Genetics. We are also very grateful to Dr. Shin-ichiro Fujio at National Museum of Japanese History and to the Center for Accelerator Mass Spectrometry at Yamagata University (YU-AMS) for their assistance with radiocarbon dating. This study was conducted in collaboration with YU-AMS and the laboratory of radiocarbon dating at the University Museum, The University of Tokyo. This work was partly supported by KAKENHI Grants-in-Aid for Scientific Research on Innovative Areas (18H05506 to M.M., 18H05511 to Y.Kawai., 18H05507 to K.S., 19H05349 and 21H00347 to R.K.), Transformative Research Areas (A) (23H04842 to R.K., 23H04844 to Y.Kawai., 23H04843 to H.K.-K., and 23H04839 to M.Takigami.), Research Activity Start-up (24K23946 to L.S.), and Scientific Research (B) (23H02566 to M.M.) from the Japan Society for the Promotion of Science (JSPS), and grant from the Wellcome Trust (224575/Z/21/Z to G.H.). This work was also supported by University of the Ryukyus (Research Project Promotion Grant for Young Researchers (22SP04102 to M.M.), the Spatiotemporal Genomics Project, and the Okinawa Bioinformation Bank Project) and by the Okinawa Prefecture (the Okinawa Innovation Ecosystem Joint Research Promotion Project for Advanced Medicine (to S.M.)).

## Author contributions

Conceptualization, M.M., H.K.-K., and R.K.; Sampling of modern samples, M.M., K.K., M.I., S.M., and R.K.; Excavating human remains, C.K., T.S., A.S., M. Takenaka., and N.D.; Sampling of ancient samples, T.K., N.A., N.D., K.S., and H.K.-K.; Radiocarbon dating, M.Takigami, M.Y., T.O., H.O., M.S., and N.K.; DNA extraction and sequencing, T.K., N.A., K.S., and H.K.-K.; Data Curation, M.M., Y.Kawai, L.S., N.B., G.H., H.K.-K., and R.K.; Analysis – Genotype calling, M.M., Y.Kawai, L.S., and H.K.-K.; Analysis – Y chromosome, Y.Kameda, and H.K.-K.; Analysis – Imputation, Y.Kawai; Analysis – F-statistics, M.M.; Analysis – treemix, M.M. and K.K.; Analysis – AncIBD and hapROH, N.B., and G.H.; Analysis – Relate, M.M., and L.S.; Analysis – fastsimcoal2, M.M.; Analysis – ALDER and RFmix, M.M.; Writing – Original Draft, M.M., Y.Kawai, M. Takigami., C. K., M.Takenaka., N.B., H.K.-K. and R.K.; Writing – Review & Editing, all members

