## Supplementary Information for "Ancient Ryukyu Jomon contributed to past and current genetic structure of Japanese populations"

### Supplementary Note 1. Archeological description of sites

#### West area of Shiitachi Site

This site is in rock shade ruins located on Gushikawa Island, which is a small island with a perimeter of 4 km located near the Izena Island. Izena Island belongs to the Okinawa Islands and is located northern west of Okinawa-jima Island. The archaeological sites of Gushikawa Island were excavated by different groups at different times: Okinawa Prefectural Board of Education in 1975, Izena Village Board of Education with Okinawa Prefectural Board of Education in 1976-1980, Izena Village Board of Education with Kagoshima University/Okinawa International University in 1989-1992, and Okinawa Prefecture Archaeological Center in 2006-2009 (Asato 1977, 1979, 1980, 1981; Kishimoto 1993; Nakayama 2012).

A mass of human bones was excavated from the third, fourth, and fifth (B) layers. Most of bones were not in anatomical order but rather were aggregated in one place in the rock shade after burial. However, several bones observed at the top of third and fourth layers were found in anatomical order. Based on excavated pottery, the bones were dated to the late Jomon, and the carbide layer near the cranial bone dated to 3,730±30BP based on C-14 radiocarbon dating (Katagiri 2012).

Drs. Matsushita and Doi conducted physical anthropological analysis of these bones. They estimated that at least 14 and 11 individuals were buried at the third/fourth and fifth (B) layer, respectively (Matsushita and Ohta 1993). Two individuals from the fifth (B) layer showed dental extractions at the lower lateral incisor (Doi 2012). These human bones are housed at the Okinawa Prefecture Archaeological Center.

#### Cliff bottom site of Gushikawa Gusuku

This site is in rock shade ruins located at the seacoast in Uruma City in the middle of the east side of Okinawa-jima Island. In 2004-2007, Dr. Doi with Gushikawa City Board of Education conducted an investigation of the site (Doi 2008). This work was supported by a KAKEN Grant-in-Aid for Scientific Research (C) titled, “Excavation at the cliff bottom site of the Gushikawa Gusuku, Okinawa”.

They excavated a mass of fragmented human bones at the third layer of this site. These human bones were not in anatomical order and included burnt bones. Since Kyushu-type Yayoi pottery was also found, the estimated date of this site is the late Yayoi. Because many shellfish products were also excavated, a fruitful shellfish culture seems to have been achieved.

Based on physical anthropological analysis of these bones, Dr. Doi estimated that these bones were from at least 67 individuals (18 male adults, 11 female adults, 19 sex unknown adults, and 19 immature individuals). Of them, bones from 31 individuals (46%) were burned, and bones from other 36 individuals (54%) were not. Failure to determine the sex of the individuals is mainly due to the bones having been burned. Several individuals had extraction of four lower incisors. These human bones are housed at the Uruma City Board of Education.

#### Hanzanbaru A Site

This site is in Kuwae, Chatan Town in the middle of the west side of Okinawa-jima Island. A Hanzanbaru colony was at the site before World War II, and after the war, this area was occupied by the U.S. Military Base (Camp Lester). In March 2015, the northern area of this base, including Hanzanbaru A Site, was returned to the Japanese government. Before its return, Chatan Town Board of Education conducted an excavation of this site in 2007-2011.

The fourth layer of this site included deposits of coastal soils. From this layer, human bones corresponding to at least 12 individuals were excavated, and were identified as Nos.01-12 in the archeological report (Shimabukuro 2016). We used human bones from Nos. 03 and 05 for ancient genome sequencing. Although these bones are organized as belonging to individuals, most individuals were missing most of the bones and had partial bones. Thus, we could not infer burial method or position of each individual.

Dr. Fujita conducted a physical anthropological analysis of these bones. Individual No. 03 was identified as a sex-unknown young adult and showed dental extractions of four lower incisors. Since there are no artifacts associated with these bones, dating is difficult. However, dental extraction from the lower jaw was common in the Ryukyu prehistoric period, suggesting that individual No. 03 is from the prehistoric period. Individual No. 05 was identified as a male adult. As for No. 03, there were no artifacts associated with these bones, and we could not estimate a date. Morphological analysis of the cranial bone showed the tendency for a small head and short face. This suggests that human bones from individual No. 05 also date to the prehistoric period (Fujita 2016). These human bones are housed at the Chatan Town Board of Education.

#### Ireibaru D Site

This site is located in Kuwae, Chatan Town, in the middle of the west side of Okinawa-jima Island. As for Hanzanbaru A Site, this site had been occupied by the U.S. Military bases (Camp Lester) after the end of World War II, and the northern area of this base was returned to the Japanese government in March 2015. The Chatan Town Board of Education conducted an excavation of this site in 2000-2001.

Five trenches (5 m × 40-100 m each) were dug for excavation. Human bones were excavated from the soil sedimentary layers in Trench 4 (5 m × 100 m), Grid 6. Although this layer is derived from the 15th or 16th centuries, the estimated date of the human bones seems to be more ancient based on the scattered positions of the bones at the excavation site, which makes it more difficult to estimate the burial method (Agarijo and Shimabukuro 2008).

Drs. Matsushita and Matsushita conducted a physical anthropological analysis of these bones. They estimated that the minimum number of individuals was three (one adult male, one adult female, and one infant), and the estimated maximum number of individuals was seven (Matsushita and Matsushita 2008). We conducted an ancient genome analysis using a bone from individual No.3, who had only the forehead bone and was identified as an adult female. These human bones are housed at the Chatan Town Board of Education.

#### Ufudoubaru Shell Mounds

This site is in rock shade ruins located in Yomitan Village in the middle of the west coast of Okinawa-jima Island. Mr. Teijun China firstly found this archeological site in 1971. At that time, most of ruins were already destroyed by sand extraction work and only a part of the rock shade had only remained.

Dr. Takamiya from Okinawa International University and his colleagues conducted excavations of this site in 1971 and 1989. They excavated human bones at Point A and Point B of the rock shade. In particular, they excavated many human bones at the lower part of first layer at Point A. These human bones were not in anatomical order but were aggregated in one place in the rock shade for burial. At the Point C, one cranium bone was excavated. This cranium bone was covered by pottery, and no other bones were recovered. These conditions suggest that only cranium bones were reburied using pottery (Takamiya et al. 1993).

Dr. Doi and her colleagues conducted a physical anthropological analysis of these bones. The estimated minimum number of individuals was 18 (12 adult males, 5 adult females, and 1 sex-unknown young adult). Since most of the human bones were derived from adults and no infant bones were excavated, burials seem to be specific to adult individuals. Three female individuals had dental extractions of four lower incisors. The estimated date based on pottery excavated along with the human bones was the late Shell-mound period (Ryukyu Jomon) (Kobashigawa et al. 2009). These human bones are housed at the Okinawa Prefecture Archaeological Center.

#### Furuzamami Shell Mounds (Siru area)

Furuzamami Shell Mounds are dune ruins located in Zamami Village on Zamami Island in the Kerama Islands. The Siru area is at the south end of this site. In 1977, human bones from five individuals were excavated in the process of sand extraction work. One individual was surrounded by rectangle-shaped plate coral containing Tridacna and showing water erosion, suggesting that this individual was buried in a cist grave.

The Alumni Association of the archeological laboratory from Okinawa International University conducted a surface exploration in 1978. The Okinawa Prefectural Board of Education conducted excavations of this site in 1981. In these investigations, scattered human bones from several individuals were excavated (Kishimoto 1982; Zamami Village History Editing Committee 1989).

Dr. Doi and her colleagues conducted a physical anthropological analysis of these bones. The estimated minimum number of individuals was seven (three adult males, three adult females, and one sex-unknown immature individual). Three individuals had dental extractions of the four lower incisors. Since no artifacts, such as pottery, were excavated from this site, dating using archeological evidence was difficult. However, dental extractions from the lower jaw are common in the Ryukyu prehistoric period, suggesting that these human bones are from the prehistoric period (Tokumine et al. 2009). These human bones are housed at the Okinawa Prefecture Archaeological Center.

#### Oike B Site

This site is at dune ruins located in Toshima Village in the Takara Island in the northern part of the Amami Islands. From this site, ruins derived from the Shell-mound period (Ryukyu Jomon) were excavated in 1993-1995. Pottery and bones of Ryukyu wild boar (*Sus scrofa riukiuanus*) were excavated at Site A. The pottery was of the Shell-mound period and two different bones of Ryukyu wild boar is dated to 4,266±22 and 4,608±24 calBP.

Well-preserved human bone (No. 1) surrounded by a cist grave was excavated at Site B. The human bones were derived from a female mature adult and showed morphological characters typical of Ryukyu prehistoric people, such as low height, small head and short face. The height was calculated to be 144.0 cm using the Pearson formula based on the length of the right femur. This individual may have been employed in physical works, such as rowing, because her deltoid of the humerus was well developed. Three bracelets made from shells (*Scutellastra optima*) were placed on her left wrist. The human temporal bone was dated to 3,165±23 calBP.

#### Tomachin Site

This site is located on Tokuno-shima Island, one of the Amami Islands. The first excavation of this site was conducted in 1992. Three sarcophagus tombs and one pit burial were excavated during the second investigation in 2004-2009. Human bones were excavated from sarcophagus tomb No. 1 (Shinzato 2013). Based on associated artifacts, this tomb was dated to the Shell-mound period, early 5 Phase (i.e., late Jomon/early Yayoi in Hondo). The tomb was separated into three sections: upper, middle, and lower rows. From the upper row, a mass of human bones not in anatomical order, including four cranial bones and three appendicular skeletons, were excavated. From the middle row, one individual was excavated. This individual was in anatomical order but without a cranial bone. Additional appendicular skeletons of other individuals were also excavated from the middle row. From the lower row, scattered human bones were observed.

Based on the positions and characters, a physical anthropological analysis of these bones was conducted (Takenaka 2013). One of the cranial bones from the upper row and bones from middle row, which were in anatomical order, seemed to be from the same individual, and we defined these bones as individual No. 4. The cranial bone of individual No. 4 may have been moved in the process of making the sarcophagus tomb for individual No. 3. The cranial bone of individual No. 4 and the appendicular skeleton of individual No. 3 were aligned with the bottom stone in the upper row. The order of burial in the upper row was individuals No. 3, No. 2, and No.1. After skeletonization of individual No. 1, the mandible, right tibia, and fibula bones were artificially moved. We utilized individual No.2 (male mature adult: Tomachin001), No. 3 (male middle-aged adult: Tomachin002), and No. 4 (female middle-aged adult: Tomachin003) for ancient genome sequencing.

#### Omonawa Shell Mounds

This site is a complex of ruins, including a shell mound, cave, and open land. The first excavation was conducted in 1982, and a box-style sarcophagus tomb was excavated from the first cave of the first shell mound (Isen Town Board of Education 1983). From this tomb, human bones from one individual (female, elderly adult) in well-preserved supine extended burial was identified. This individual showed typical morphological characters of Ryukyu prehistoric people, such as low height, small head and short face.

In 2013, a second excavation of the first shell mound was conducted, and human bones were excavated from the fourth and fifth layer of Trench C. From the east side of the fourth layer of Trench C, only a cranial bone was excavated (Isen Town Board of Education. 2016). Based on morphological analysis, this cranial bone is derived from a male, matured adult who had a small head and low orbit placement with a deeply recessed root of the nose. The bone was dated to 896-807 BC (2,845-2,756 BP, 2σ). This corresponds to the Shell-mound period, early 5 Phase (i.e., late Jomon/early Yayoi in Hondo). Limestone clast the size of a human head, small-rounded pebbles, nail-marked pottery, and scattered human bones were excavated from the fifth layer of Trench C. We conducted ancient genome sequencing on human bone from the fifth layer of Trench C (Omonawa002).

#### Gushibaru Shell Mounds

This site is in dune ruins located at the southern coast of Ie Island, which is in the western part of the Okinawa Islands. This site was excavated by different groups at different times: Dr Tomoyori in 1963, Ie Town Board of Education with the Okinawa Prefectural Board of Education in 1984, and Okinawa Prefectural Board of Education in 1995 (Tomoyori 1968, 1970; Asato 1985; Kishimoto 1997). Signatures of trade between Ryukyu and Kyushu (south part of Hondo) were identified among the artifacts from the most recent excavation (Kishimoto 1997). For example, Yayoi pottery from Kyushu was firstly excavated in Okinawa Prefecture and accumulated remains of shells, such as *Sinustrombus latissimus* and Cone snails, which are always associated with Yayoi pottery through the Japanese Archipelago were excavated from this site.

Human bones were excavated from the fifth layer of Trench 1, south area and the seventh layer of Trench 2, south area. Human bones from the fifth layer of Trench 1 did not include cranial bone and were not in anatomical order. Since there are no artifacts associated with this bone, we could not provide a date. However, sedimentation status of the dune ruins suggests that these are from the prehistoric period.

Although human bones from the seventh layer of Trench 2 were scattered, the number of appendicular skeletons suggests that these bones are from several individuals. The associated pottery was dated to the late Shell-mount period (i.e., late Yayoi-Heian in Hondo). In particular, 86 perforated ducal thorny oysters associated with human bones were excavated. This is a very rare finding (Kishimoto 1985). This site was registered as a national historic site in Japan, and the human bones are housed at the Okinawa Prefecture Archaeological Center.

#### Tsuken Shell Mounds

This site is in dune ruins located at Tsuken Island (Tsuken, Uruma City, Okinawa Prefecture), which is near the east side of Okinawa-jima Island and belongs to Okinawa Islands. This small island has a perimeter of 7.6 km. In the process of construction work, human bones from two individuals were discovered in 1974. This is the first finding of this site. Then, Okinawa Prefectural Board of Education conducted an excavation in the same year. At that time, interviews of residents provided the information that human bones had already been excavated in 1920 (Kin et al. 1975).

Human bones were excavated from white sand layer, Area 3, which is approximately 120 m below the ground surface. Although this layer looks white, soils surrounding human bones are dark brown in color. The human bones were positioned supine with the head toward the north for burial. Human bones excavated at the construction site had the same burial style, and individuals were spaced about 2 m apart. Bones from all four human individuals (one excavated in 1920, two excavated in 1974, and one excavated in later) were buried in a line (Toma 1975).

Since no artifacts associated with human bones were excavated, it is difficult to estimate the date for these individuals. However, pottery dating to the late Shell-mound period (i.e., late Yayoi-Heian in Hondo) was excavated from a layer lower than the white sand layer. This suggests that these human bones are from at least after the Yayoi period in Hondo. These human bones are housed at the Okinawa Prefecture Archaeological Center.

#### Shiraho Saonetabaru Cave Ruins

This site is a cave ruin located on the east coast of Ishigaki Island belonging to Yaeyama Islands. This site was discovered in 2008, and Paleolithic human bones were excavated in 2009 (Nakagawa et al. 2010). Because of this finding, a large-scale excavation was conducted by the Okinawa Prefecture Archaeological Center collaborating with Ishigaki City Board of Education and the Okinawa Prefectural Museum in 2010 and 2012-2016 (Nakaza 2013, 2017a, 2017b; Katagiri 2019).

This site seems to have been continuously used from the Paleolithic to the Nakamori period of Ishigaki Island (same time as the Gusuku period in Ryukyu). The first finding of mass of Paleolithic human bones on Ishigaki Island highlights the importance of this site. One of the other prominent findings is that human bones derived from the Shimotabaru period (same time as the Shell-mound period in Ryukyu, but different culture) were first excavated on Ishigaki Island. Human bones derived from the Shimotabaru period were only used for ancient genome sequencing in this study.

Human bones from the Shimotabaru period were excavated from the Layer S, and they were not kept anatomical order but were aggregated near the wall after burial. Although no pottery was excavated, carbide near the human bones was dated to 4,090±30BP and direct dating of the human bone placed it at 3,970±30BP.

Drs. Doi, Kono and their colleagues conducted a physical anthropological analysis of these bones. They estimated that these bones include at least 3 individuals (1 male adult, 1 female adult, and 1 immature individual) (Doi et al. 2017). This site was registered as a national historic site in Japan, and the human bones are housed at the Okinawa Prefecture Archaeological Center.

### Supplementary Note 2. C14 dating of ancient bone

Chunk or powder bone samples (200-500mg) were collected from skull bones. In the case of chunk bone samples, surface contaminations such as soil and adhesive were physically removed with a dental drill. After soaking the samples in acetone for 10 min, we conducted ultrasonic cleaning in ultrapure water for 10 min. Bone powder was sampled from internal area of bone, not adhered with soil, together with samples for DNA analysis. After removing carbonate using 0.6 M HCl for 24 h, samples were soaked in up to 0.2 M NaOH for a few hours to remove exogenous organic matters such as fumic acid. Then, collagen was extracted by heating in weak hydrochloric acid (pH 4) at 90°C for 24 h. The solution was filtered through a glass filter, which had a particle retention capacity of 0.7 μm, and then freeze-dried. The collagen samples were sent to the Center for Accelerator Mass Spectrometry (AMS) at Yamagata University, and carbon and nitrogen isotope analysis as well as radiocarbon dating were conducted. Because only a small amount of bone powder could be collected from the sample of Gushikawa002, micro-measurement of radiocarbon was conducted at the University Museum, The University of Tokyo.

Collagen preservation was estimated from the collagen extraction rate, carbon and nitrogen content and their molar ratio (DeNiro 1985; van Klinken 1999). Marine resource consumption was calculated with the ISOCONC1.01 program using carbon and nitrogen isotope ratios (Phillips and Koch 2002). Radiocarbon ages were calibrated with the Oxcal 4.4.4 program (Bronk Ramsey 2009) using the mixed curve of IntCal20 and Maine20 with the contribution rate for marine resources described above (Heaton et al. 2020; Reimer et al. 2020). The local reservoir effect (ΔR value) was estimated to be -143±33, which is corrected to the latest calibration curve referenced from the CALIB.org program (Stuiver and Reimer 1993; Yoneda et al. 2007).

### Supplementary Note 3. Ancient DNA extraction and library preparation

The temporal bones/teeth of human skeletons (Supplementary Note 1) were subjected to DNA analysis. Collection of bone powder and DNA extraction were performed on a bench in a clean room at the University of Yamanashi. Clean suits, masks, and caps, as well as double gloves, were worn during the work. The temporal bones were drilled from the top of the petrous region, which is pyramidal shaped, including the inner ear. Then, approximately 200 mg of bone powders was collected. The root of the teeth was cut, and the dental pulp cavities were drilled. The tooth roots were crushed to obtain bone powder. DNA extraction was performed according to the method of Adachi et al. (2013).

Maximally four DNA libraries were constructed for each sample to sequence whole genomes. One DNA library was constructed by modifying the "no uracil-DNA-glycosylase treatment" method of Rohland et al. (2015). The remaining libraries were created using a partially modified version of the "half uracil-DNA-glycosylase treatment" method of Rohland et al. (2015). These libraries were used to exclude deamination. The DNA libraries were prepared in a clean room at the National Museum of Nature and Science. In total, maximally four DNA libraries were amplified twice by PCR to ensure the supply of DNA required for target enrichment of whole genomic DNA, if necessary. For enrichment, MYbaits WGE (Daicel Arbor Biosciences) was used, working according to protocol and with the hybridization temperature at 55°C.

### Supplementary Note 4. NCBN Controls WGS Consortium

Hatsue Ishibashi-Ueda^1^, Tsutomu Tomita^1^, Michio Noguchi^1^, Ayako Takahashi^1^, Yu-ichi Goto^2^, Sumiko Yoshida^3^, Kotaro Hattori^3^, Ryo Matsumura^3^, Aritoshi Iida^4^, Yutaka Maruoka^5^, Hiroyuki Gatanaga^6^, Akihiko Shimomura^5^, Masaya Sugiyama^7^, Satoshi Suzuki^5^, Kengo Miyo^8^, Yoichi Matsubara^9^, Akihiro Umezawa^10^, Kenichiro Hata^11^, Tadashi Kaname^12^, Kouichi Ozaki^13^, Haruhiko Tokuda^13^, Hiroshi Watanabe^13^, Shumpei Niida^13^, Eisei Noiri^14^, Koji Kitajima^14^, Yosuke Omae^14,15^, Reiko Miyahara^14^, Hideyuki Shimanuki^14^, Yosuke Kawai^15^, and Katsushi Tokunaga^14,15^

^1^NCVC Biobank, National Cerebral and Cardiovascular Center, Suita, Osaka 564-8565, Japan

^2^Medical Genome Center, National Center of Neurology and Psychiatry, Kodaira, Tokyo 187-8551, Japan

^3^Department of Bioresources, Medical Genome Center, National Center of Neurology and Psychiatry, Kodaira, Tokyo 187-8551, Japan

^4^Department of Clinical Genome Analysis, Medical Genome Center, National Center of Neurology and Psychiatry, Kodaira, Tokyo 187-8551, Japan

^5^NCGM Biobank, National Center for Global Health and Medicine, Shinjuku-ku, Tokyo 162-8655, Japan

^6^AIDS Clinical Center, National Center for Global Health and Medicine, Shinjuku-ku, Tokyo 162-8655, Japan

^7^Department of Viral Pathogenesis and Controls, Research Institute, National Center for Global Health and Medicine, Ichikawa, Chiba 272-8516, Japan

^8^Center for Medical Informatics and Intelligence, National Center for Global Health and Medicine, Shinjuku-ku, Tokyo 162-8655, Japan

^9^National Center for Child Health and Development, Setagaya-ku, Tokyo 157-8535, Japan

^10^Center for Regenerative Medicine, National Center for Child Health and Development, Setagaya-ku, Tokyo 157-8535, Japan

^11^Department of Maternal-Fetal Biology, National Center for Child Health and Development, Setagaya-ku, Tokyo 157-8535, Japan

^12^Department of Genome Medicine, National Center for Child Health and Development, Setagaya-ku, Tokyo 157-8535, Japan

^13^Research Institute, National Center for Geriatrics and Gerontology, Obu, Aichi 474-8511, Japan

^14^Central Biobank, National Center Biobank Network, Shinjuku-ku, Tokyo 162-8655, Japan

^15^Genome Medical Science Project (Toyama), Research Institute, National Center for Global Health and Medicine, Shinjuku-ku, Tokyo 162-8655, Japan

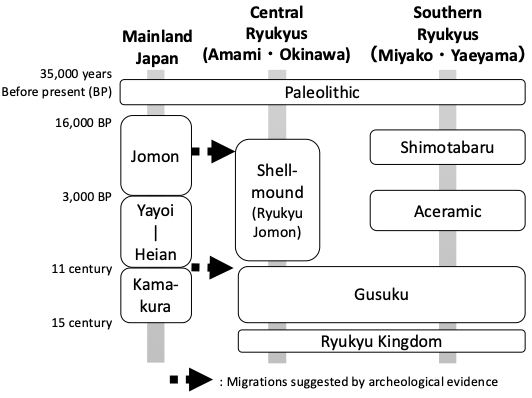

### Supplementary Fig. 1: Historical periods of Ryukyu and Hondo

The Northern Ryukyus basically follow the same chronology as the Hondo region.

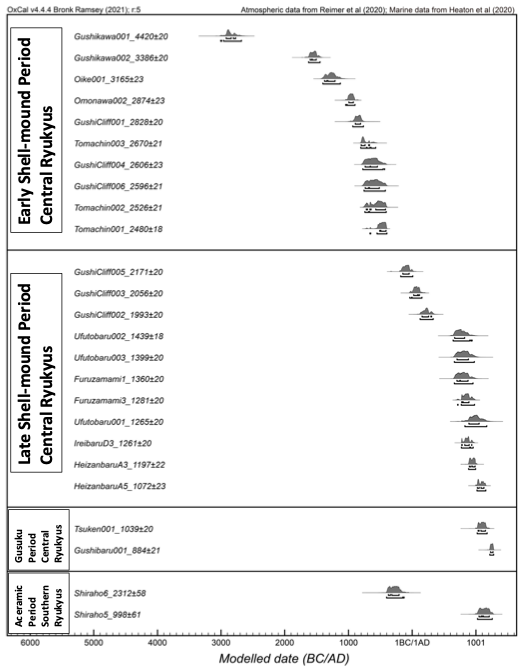

### Supplementary Fig. 2: Radiocarbon dating for Ryukyu ancient samples

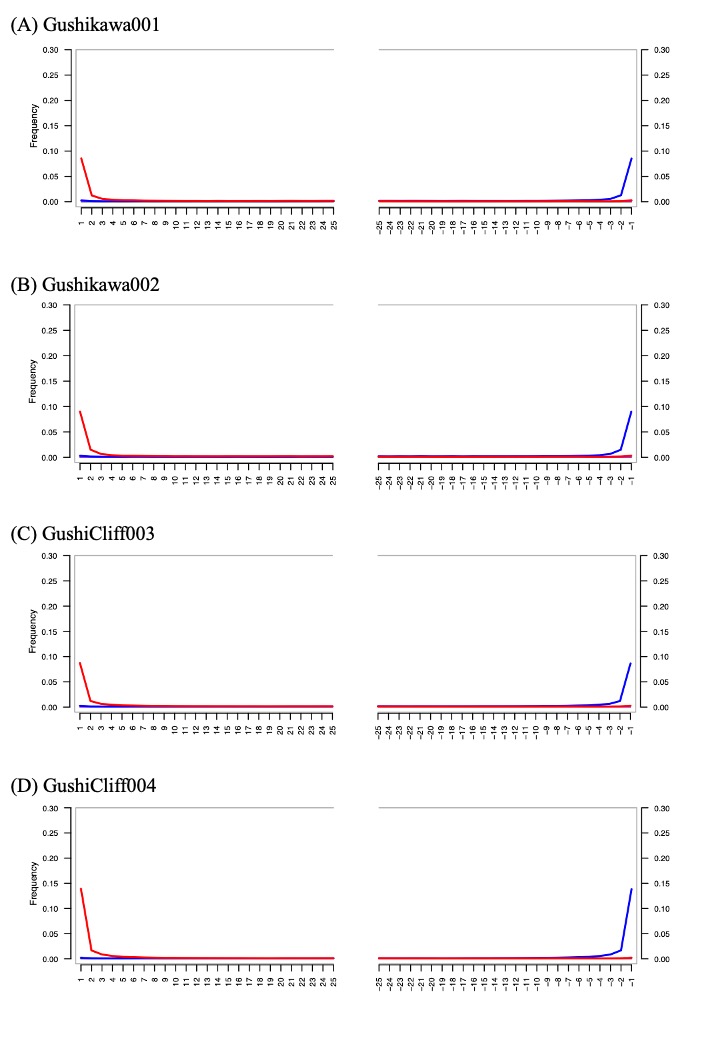

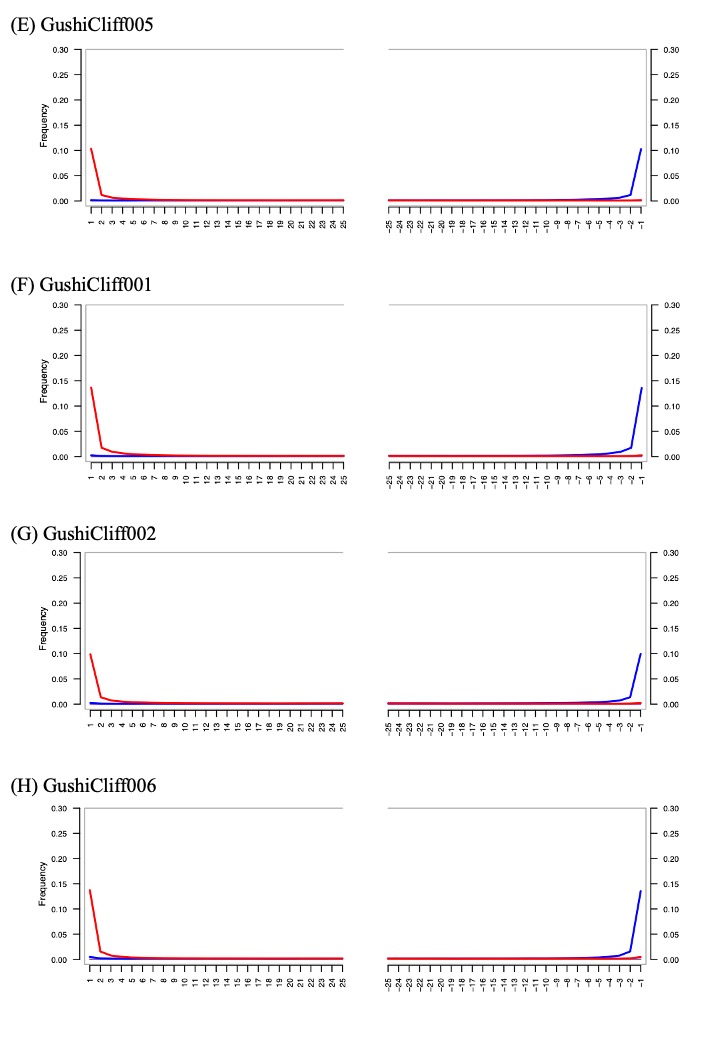

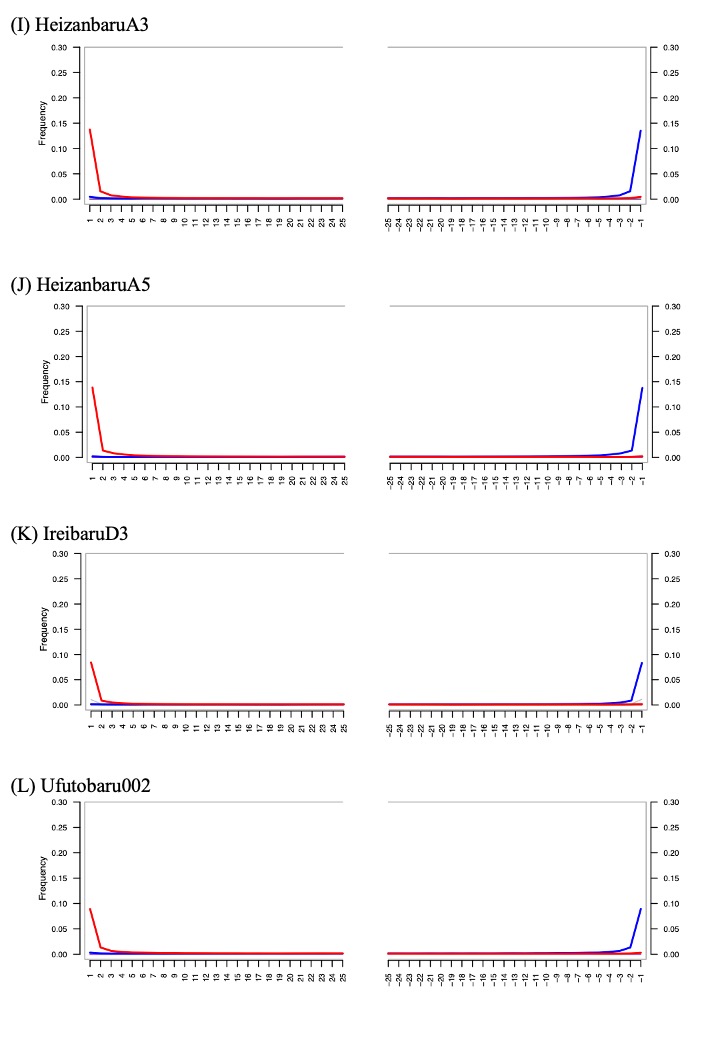

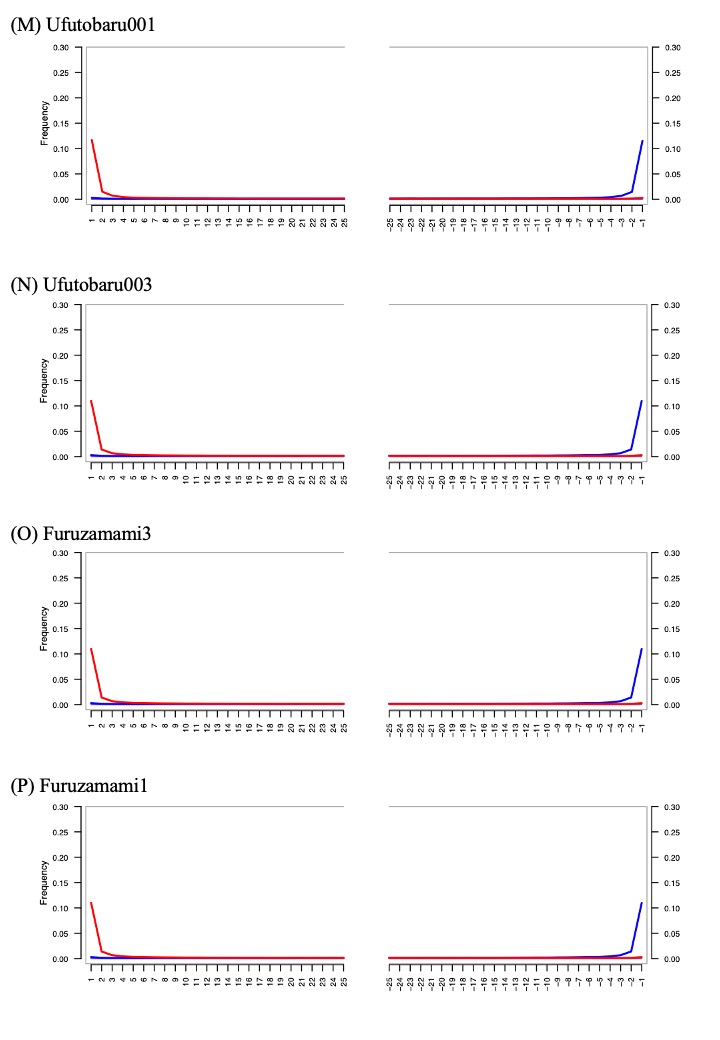

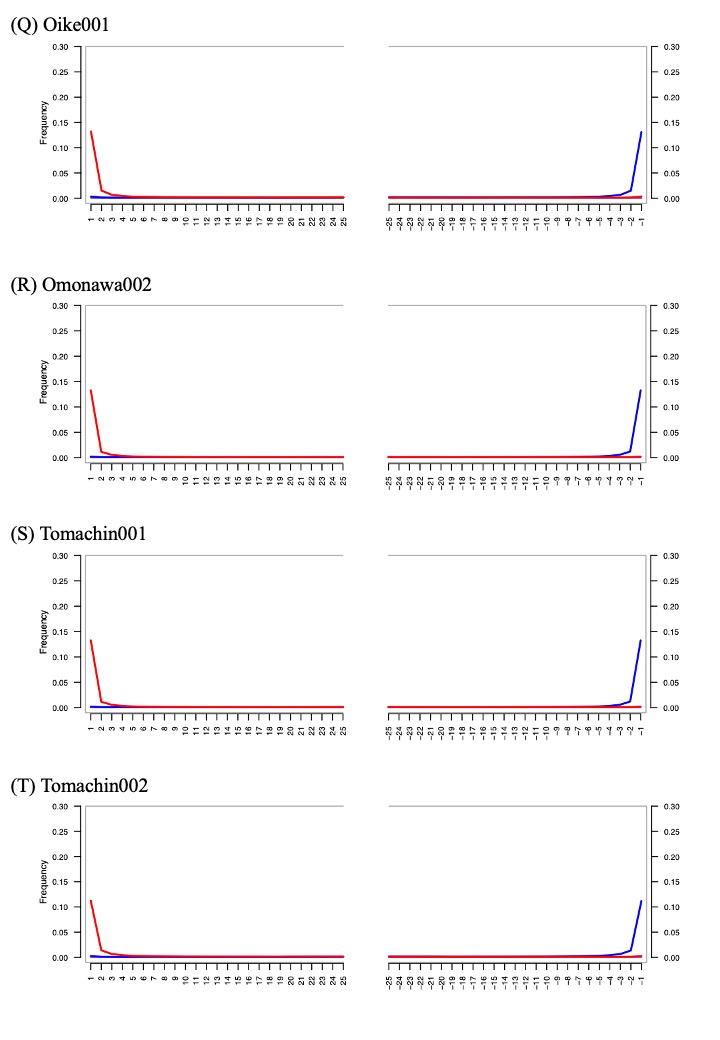

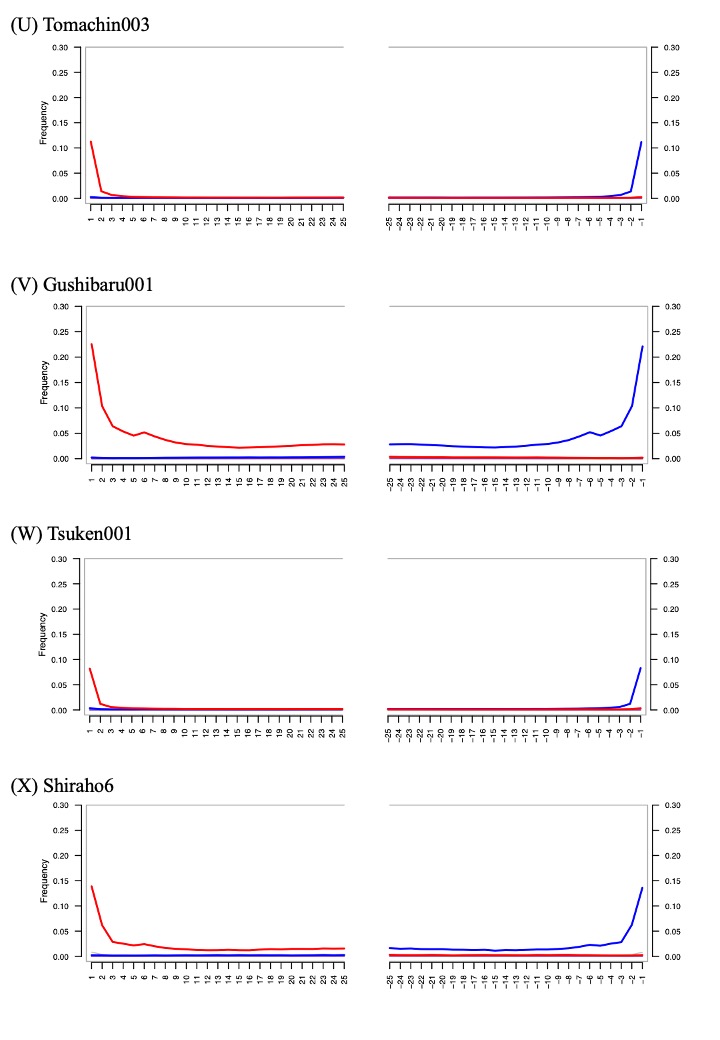

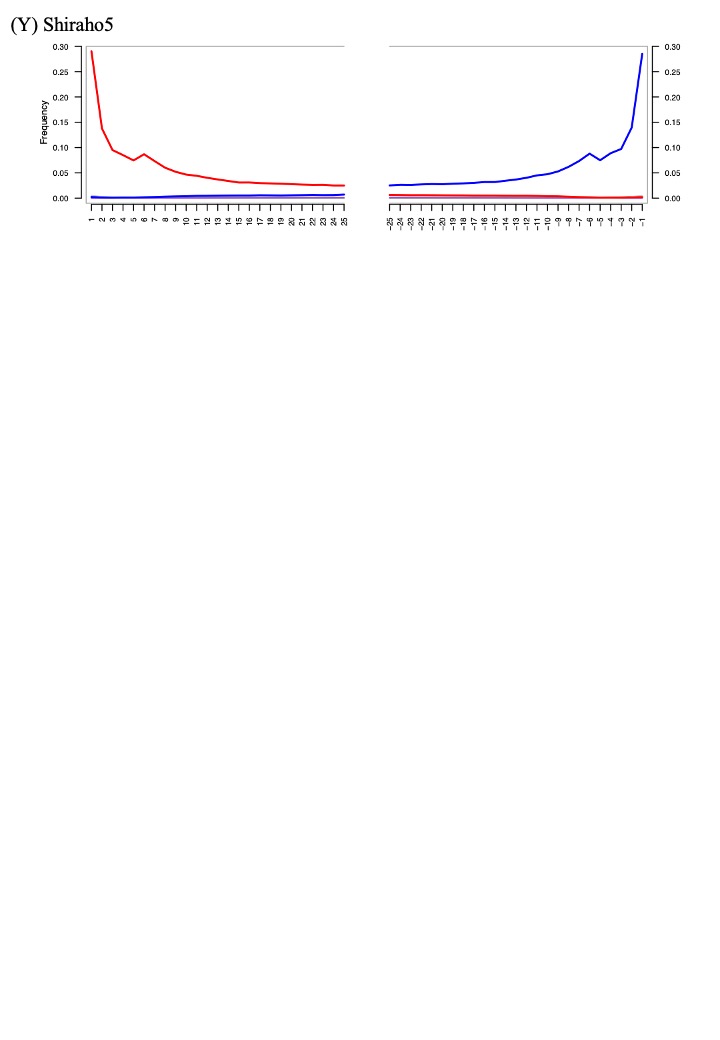

### Supplementary Fig. 3: Pattern of postmortem misincorporation

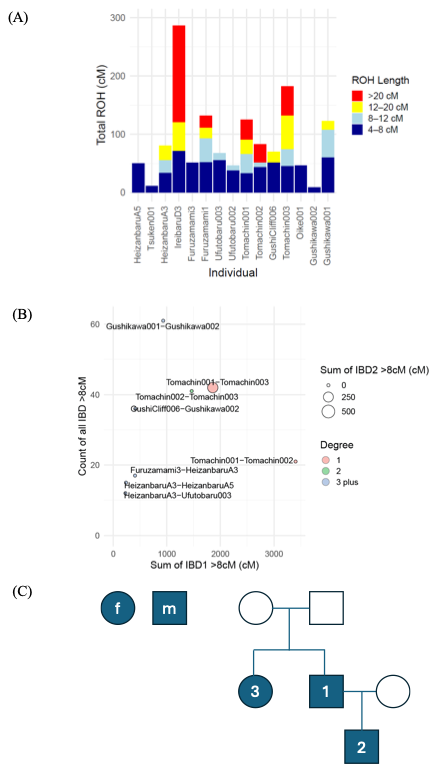

### Supplementary Fig. 4: Genomic sharing between newly sequenced individuals

**(A)** Total run of homozygosity (ROH) (cM) in each ancient individual, split into different sequence length categories. Individuals are ordered by age, based on radiocarbon dating. **(B)** Identity by descent (IBD) segment sharing between pairs of ancient individuals. Plot of sum of IBD1 segments >8cM against count of any IBD segments >8cM. Point size indicates sum of IBD2 segments >8cM. **(C)** Proposed family tree for the Tomachin individuals. We estimated that Tomachin003 and Tomachin001 were siblings. However, the genomic regions in which two IBD segments were shared between Tomachin001 and Tomachin003 accounted for only ~10% of their genome, compared with an expected 25% for full siblings, suggesting a more complicated relationship or the possibility that some IBD segments were not detected by this method.

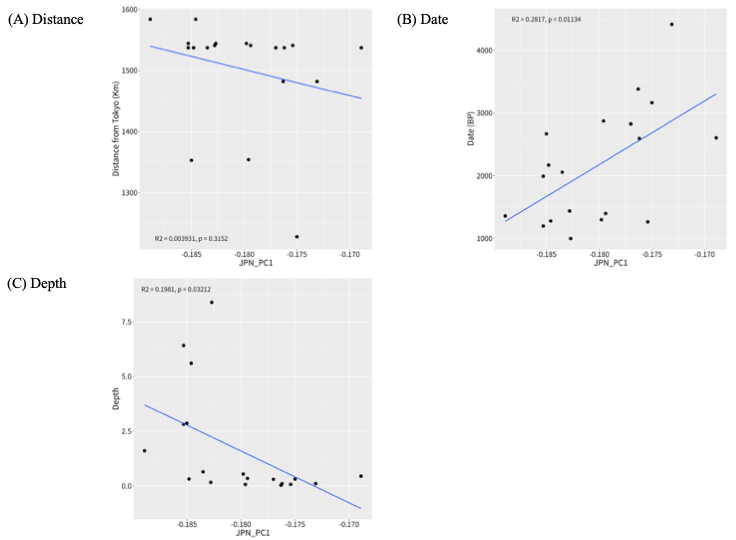

### Supplementary Fig. 5: Scatterplot between PC1 and other parameters

PC1 values of Ryukyu Jomon obtained by PCA using only Japanese populations (Fig. 1D) were plotted against distance from Tokyo (A), date (B), and sequencing depth (C). Regression lines were also plotted.

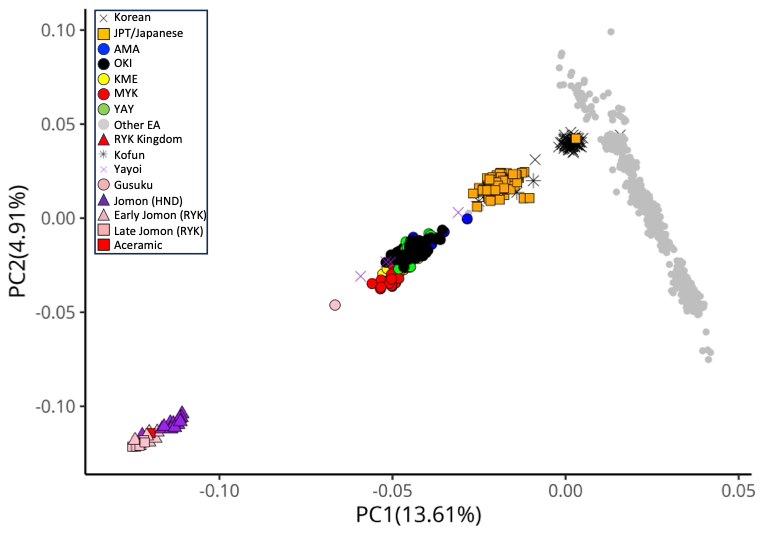

### Supplementary Fig. 6: PCA plot using imputed East Asian populations

Imputed genome sequences were used for PCA. After imputation, we filtered out variants showing minor allele frequency < 0.01, and the remaining 10,919,890 SNPs were used for PC calculation.

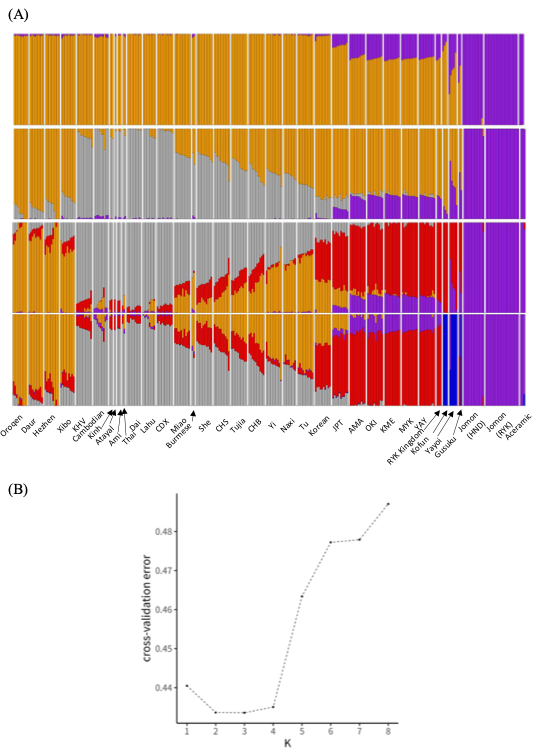

### Supplementary Fig. 7: Admixture plots with different K values

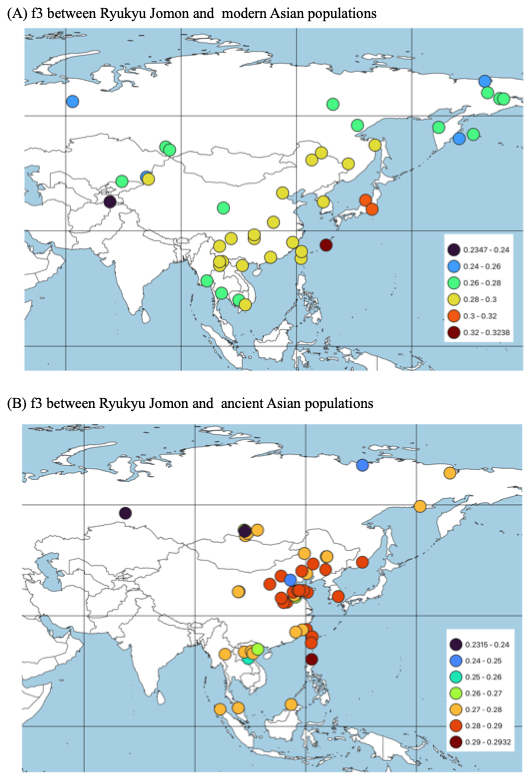

### Supplementary Fig. 8: Distribution of outgroup f3 statistics between Ryukyu Jomon and surrounding modern and ancient populations

F3 statistics were calculated using Mbuti as outgroup. Details of the f3 calculation are given in Supplementary Tables 5 and 6.

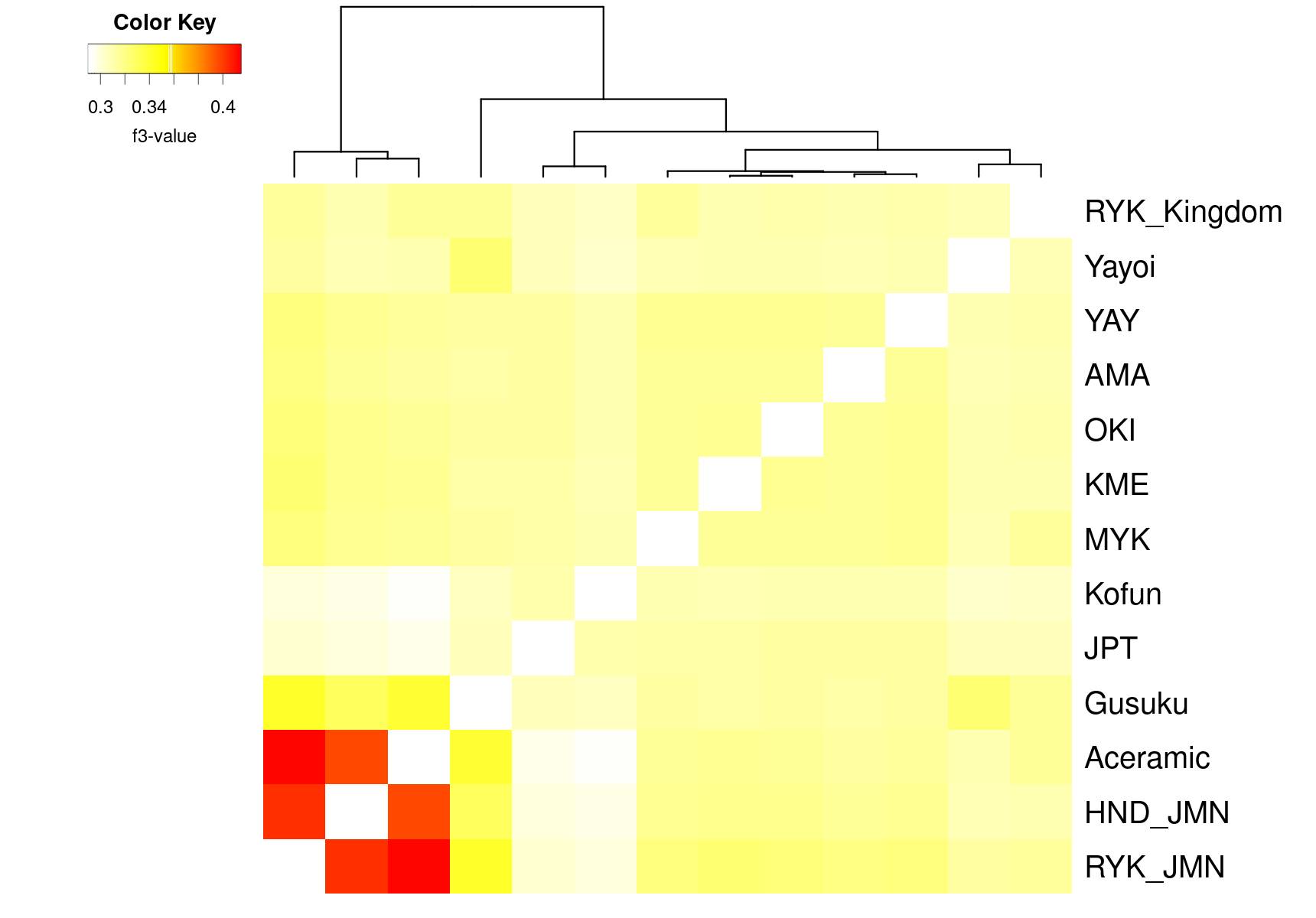

### Supplementary Fig. 9: Heatmap of population-level outgroup f3 statistics among Japanese populations

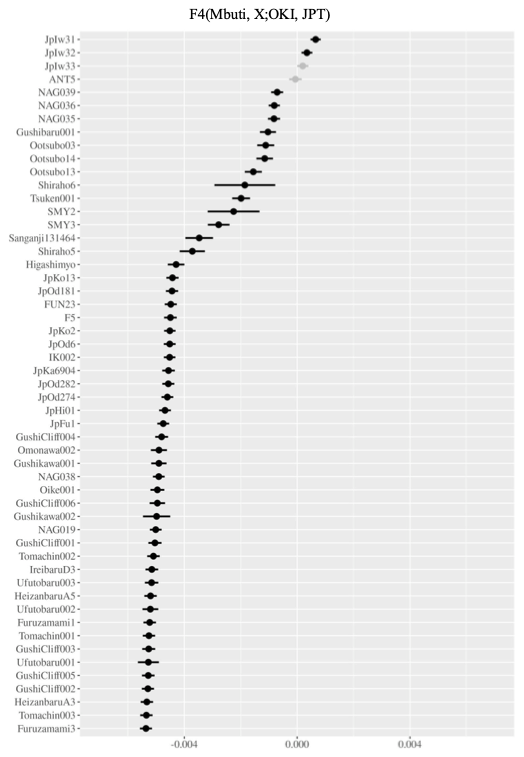

### Supplementary Fig. 10: Individual-level f4 statistics inferring admixture from ancient to modern Japanese

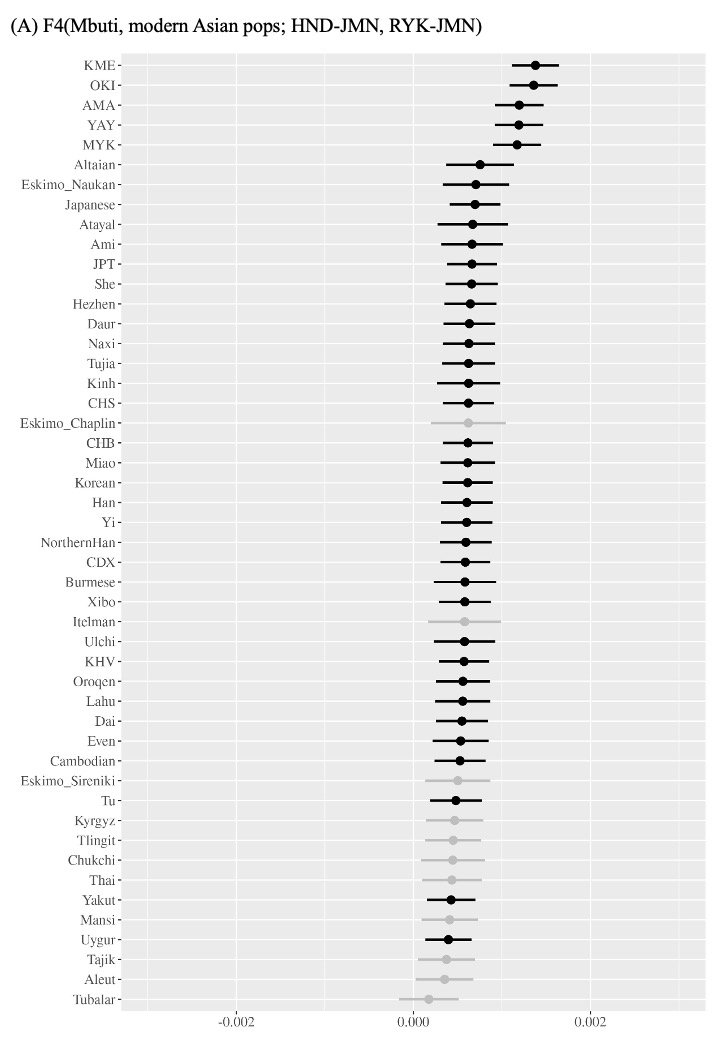

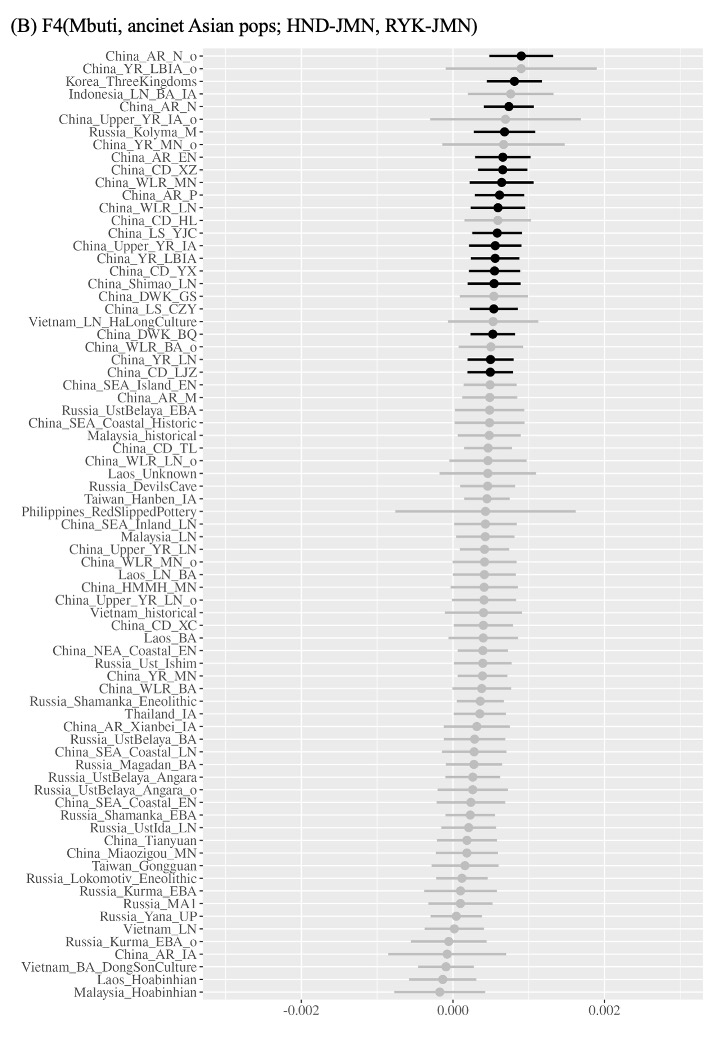

### Supplementary Fig. 11: f4 statistics using Ryukyu and Hondo Jomon as targets

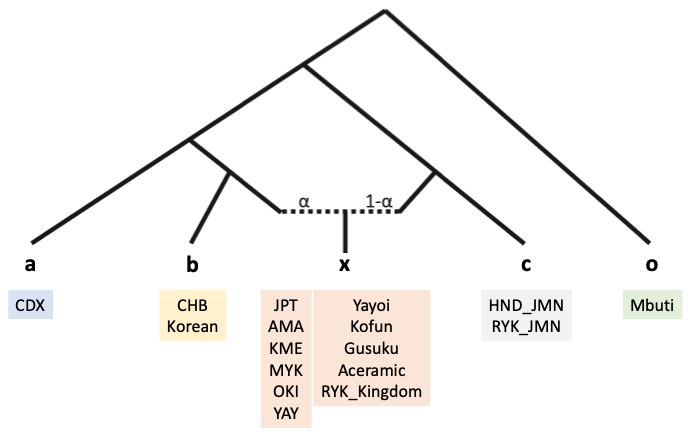

### Supplementary Fig. 12: Schematic of f4 ratio test combinations

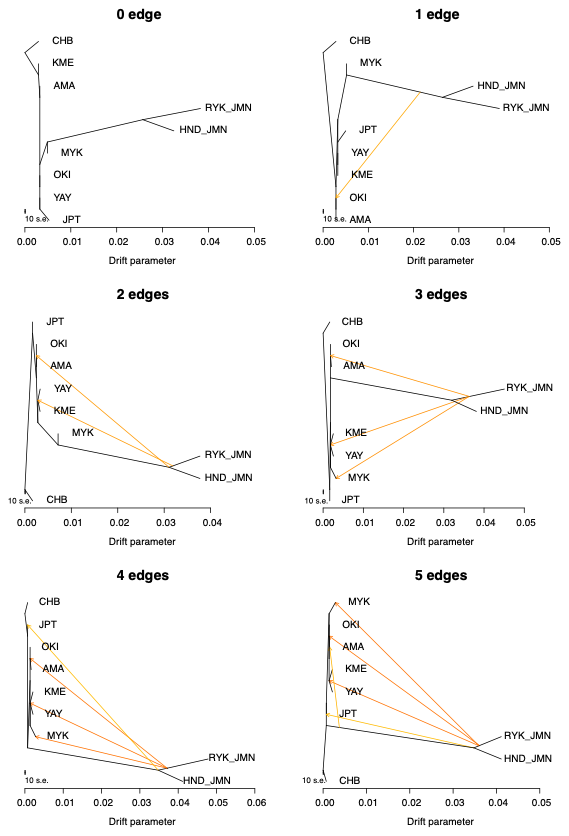

### Supplementary Fig. 13: Maximum likelihood trees using ancient and modern Japanese populations

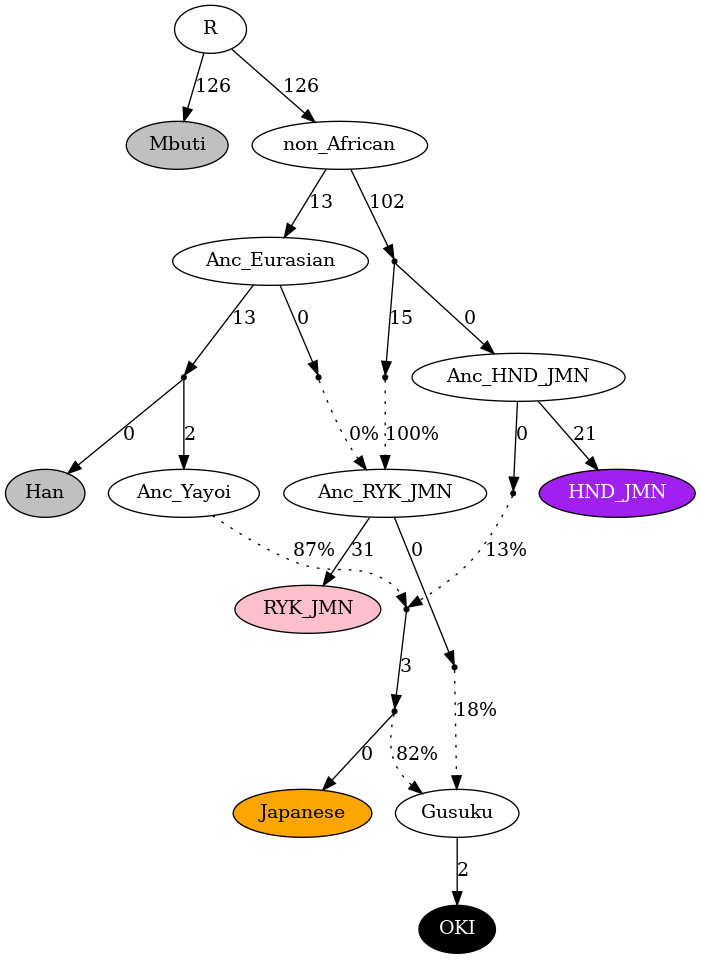

### Supplementary Fig. 14: Admixture modeling for Ryukyu populations considering admixture from ancestral Eurasian into Ryukyu Jomon

In this model, two ancestral Jomon populations were assumed, and these populations were split from an ancestral non-African population. The ancestral Ryukyu Jomon population also received admixture from an ancestral Eurasian population. Colored nodes represent actual populations as input data, while others are hypothetical populations.

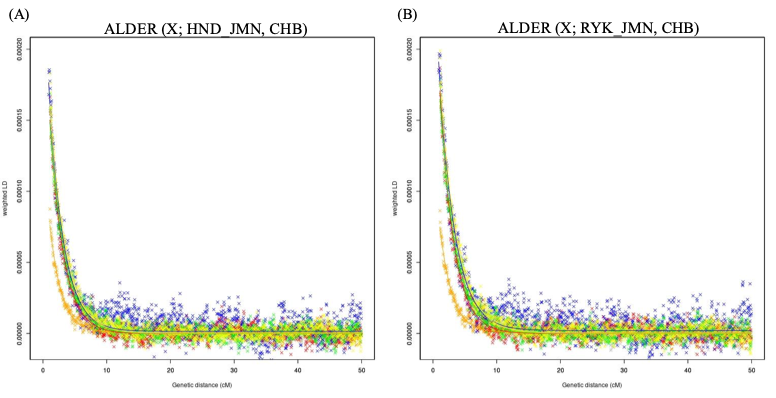

### Supplementary Fig. 15: Admixture dating of Jomon ancestry

LD decay curves for modern Japanese using CHB and Hondo Jomon (A) or Ryukyu Jomon (B) as reference. Colors are the same as in Fig. 1 and represents each Japanese populations; JPT (yellow), AMA (blue), OKI (black), KME (orange), MYK (red), and YAY (green).

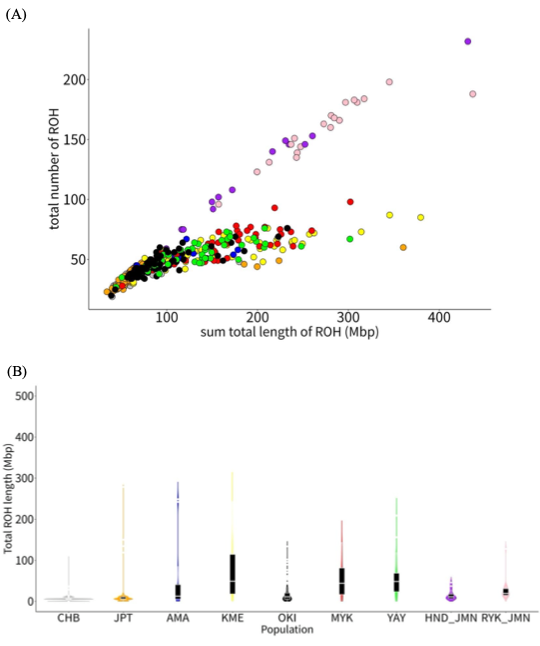

### Supplementary Fig. 16: Distribution of ROH number and length

(A) Scatter plot between total number of ROH and total length of ROH. (B) Violin plot of long ROH (more than 5 Mbp) for each population. Colors are the same as in Fig. 4.

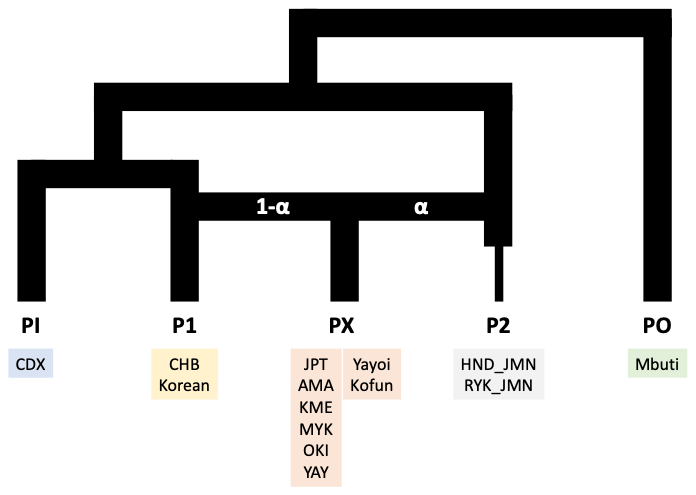

### Supplementary Fig. 17: Schematic of the twigstats test

de Barros Damgaard P, Martiniano R, Kamm J, Moreno-Mayar JV, Kroonen G, Peyrot M, Barjamovic G, Rasmussen S, Zacho C, Baimukhanov N, Zaibert V, Merz V, Biddanda A, Merz I, Loman V, Evdokimov V, Usmanova E, Hemphill B, Seguin-Orlando A, Yediay FE, Ullah I, Sjögren KG, Iversen KH, Choin J, de la Fuente C, Ilardo M, Schroeder H, Moiseyev V, Gromov A, Polyakov A, Omura S, Senyurt SY, Ahmad H, McKenzie C, Margaryan A, Hameed A, Samad A, Gul N, Khokhar MH, Goriunova OI, Bazaliiskii VI, Novembre J, Weber AW, Orlando L, Allentoft ME, Nielsen R, Kristiansen K, Sikora M, Outram AK, Durbin R, Willerslev E. 2018. The first horse herders and the impact of early Bronze Age steppe expansions into Asia. *Science.* 360:eaar7711.

DeNiro MJ. 1985. Postmortem preservation and alteration of in vivo bone collagen isotope ratios in relation to palaeodietary reconstruction. *Nature* 317: 806-809.

Doi N (Ed.). 2008. Excavation at the cliff bottom site of the Gushikawa Gusuku, Okinawa: understanding for the transition from Ryukyu prehistoric to Gusuku period. *Published by Author.* 117 pp. (in Japanese).

Doi N. 2012. Chapter 11: Human bones at Archaeological sites of Gushikawa island. In Archaeological sites of Gushikawa island (pp.245-260). *Okinawa Prefecture Archaeological Center.* (in Japanese).

Doi N, Tokumine R, Katagiri C, Kono R. 2017. Analysis of Excavated human bones (human bones). In Shiraho Saonetabaru Cave Ruins: Second report on the survey to confirm the extent of important archeological site –Summary report edition– (pp. 64-85). *Okinawa Prefecture Archaeological Center.* (in Japanese).

Fu Q, Li H, Moorjani P, Jay F, Slepchenko SM, Bondarev AA, Johnson PL, Aximu-Petri A, Prüfer K, de Filippo C, Meyer M, Zwyns N, Salazar-García DC, Kuzmin YV, Keates SG, Kosintsev PA, Razhev DI, Richards MP, Peristov NV, Lachmann M, Douka K, Higham TF, Slatkin M, Hublin JJ, Reich D, Kelso J, Viola TB, Pääbo S. 2014. Genome sequence of a 45,000-year-old modern human from western Siberia. *Nature*. 514:445-9.

Fujita Y. 2016. Chapter 4, third section: human bones excavated from Hanzanbaru A Site. In Hanzanbaru A Site (pp.424-438). *Chatan Town Board of Education.* (in Japanese).

Gakuhari T, Nakagome S, Rasmussen S, Allentoft ME, Sato T, Korneliussen T, Chuinneagáin BN, Matsumae H, Koganebuchi K, Schmidt R, Mizushima S, Kondo O, Shigehara N, Yoneda M, Kimura R, Ishida H, Masuyama T, Yamada Y, Tajima A, Shibata H, Toyoda A, Tsurumoto T, Wakebe T, Shitara H, Hanihara T, Willerslev E, Sikora M, Oota H. 2020. Ancient Jomon genome sequence analysis sheds light on migration patterns of early East Asian populations. *Commun Biol.* 3:437.

Gelabert P, Blazyte A, Chang Y, Fernandes DM, Jeon S, Hong JG, Yoon J, Ko Y, Oberreiter V, Cheronet O, Özdoğan KT, Sawyer S, Yang S, Greytak EM, Choi H, Kim J, Kim JI, Jeong C, Bae K, Bhak J, Pinhasi R. 2022. Northeastern Asian and Jomon-related genetic structure in the Three Kingdoms period of Gimhae, Korea. *Curr Biol.* 32:3232-3244.e6.

Heaton T, Köhler P, Butzin M, Bard E, Reimer R, Austin W, Bronk Ramsey C, Grootes P, Hughen K, Kromer B, Reimer P, Adkins J, Burke A, Cook M, Olsen J, Skinner L. 2020. Marine20 - the marine radiocarbon age calibration curve (0–55,000 cal BP). *Radiocarbon*. 62: 779-820.

Ie village Board of Education. 2020. In-village site excavation report. *Report on the Survey of Cultural Properties in Ie Village.* 16:62-68 (In Japanese).

Isen Town Board of Education. 1983. Excavation Report of Isen Town (1): Omonawa First and Second Shell Mounds. *Isen Town Board of Education.* 80 pp. (in Japanese).

Isen Town Board of Education. 2016. Excavation Report of Isen Town (16): Summary report of Omonawa site. *Isen Town Board of Education.* 222 pp. (in Japanese)

Kanzawa-Kiriyama H, Kryukov K, Jinam TA, Hosomichi K, Saso A, Suwa G, Ueda S, Yoneda M, Tajima A, Shinoda KI, Inoue I, Saitou N. 2017. A partial nuclear genome of the Jomons who lived 3000 years ago in Fukushima, Japan. *J Hum Genet.* 62:213-221.

Kanzawa-Kiriyama H, Kameda Y, Kakuda T, Adachi N, Shinoda K. 2023. Nuclear Genome Analysis of Human Skeletons Excavated from Otubo Shell Mound in the End of Late Yayoi, Uki-city, Kumamoto Pref. *Bull. Nat. Mus. Japan*. 242: 123–132 (in Japanese).

Katagiri C. 2012. Burials and Burial methods for prehistoric people at Gushikawa island. In the 2012 Research Symposium of Okinawa Archaeological Society: Burials and Burial methods for prehistoric period (pp.41-52). *Okinawa Archaeological Society.* (in Japanese).

Katagiri C (Ed.). 2019. Shiraho Saonetabaru Cave Ruins: Third report on the survey to confirm the extent of important archeological site –Supplementary edition–. *Okinawa Prefecture Archaeological Center*. (in Japanese).

Kin S, Asato S, Toma T. 1975. Emergency Excavation Report of Tsuken Second Shell Mounds. In Annual Report of Cultural Properties for Fiscal Year Showa 49 (pp.49-74). *Okinawa Prefectural Board of Education.* (in Japanese).

Kinoshita N, Sakamoto M, Takigami M. 2020a. Archaeological Report on the Chronology of Human Bones and Shell Accumulations of the Shell Midden Period Excavated in Uruma, Okinawa. *Bull. Nat. Mus. Japan*. 219:301-312 (in Japanese).

Kinoshita N, Sakamoto M, Takigami M. 2020b. Archaeological Report on the Chronology of Human Bone of Early Shell Midden Period etc. Excavated at Oike Site B, Takarajima, Kagoshima. *Bull. Nat. Mus. Japan*. 219:231-241 (in Japanese).

Kinoshita N, Sakamoto M, Takigami M. 2021. Archaeological Report on the Chronology of Human Bones of the Late Shell mound Period Excavated in Okinawa Islands: Gushikawajima Site Group, Gushibaru Shell mound, Gushiken Shell mound, Uhutobaru Shell mound and Gushikawa-Gusuku Under Cliff Site. *Bull. Nat. Mus. Japan*. 229:247-277 (in Japanese).

Kishimoto Y (Ed.). 1982. Furuzamami Shell Mounds: Report on the survey to confirm the extent of archeological site. *Okinawa Prefectural Board of Education.* (in Japanese).

Kishimoto Y. 1985. Ruins. In Summary of Gushibaru Shell Mound at Ie island (pp.13-19). *Okinawa Prefectural Board of Education*. (in Japanese).

Kishimoto Y (Ed.). 1993. Archaeological sites of Gushikawa island. *Izena village Board of Education*. (in Japanese).

Kishimoto Y (Ed.). 1997. Excavation Report for Gushibaru Shell Mound. *Okinawa Prefectural Board of Education.* (in Japanese).

Kobashigawa T, Katagiri C, Tokumine R, Motomura M, Oshiro A, Tengan M, Sugawara H, Doi N, Yoneda M. 2009. Human Remains recovered from the Ufutobaru Shellmidden, Yomitan Village, Okinawa Prefecture -Physical characteristics of the Ufutobaru population -. *Bulletin of the Archaeological study of Okinawa.* 6:27-40 (in Japanese).

Mao X, Zhang H, Qiao S, Liu Y, Chang F, Xie P, Zhang M, Wang T, Li M, Cao P, Yang R, Liu F, Dai Q, Feng X, Ping W, Lei C, Olsen JW, Bennett EA, Fu Q. 2021 The deep population history of northern East Asia from the Late Pleistocene to the Holocene. *Cell*. 184:3256-3266.e13.

Matsushita T, and Ohta J. 1993. Appendix: Ancient human bones excavated from archaeological sites of Gushikawa island. In Archaeological sites of Gushikawa island (pp. 215-244). *Izena village Board of Education.* (in Japanese).

Matsushita T, and Matsushita M. 2008. Chapter 6: Human bone excavated from Ireibaru D Site (1). In Ireibaru D Site (pp. 212-232). *Chatan Town Board of Education.* (in Japanese).

McColl H, Racimo F, Vinner L, Demeter F, Gakuhari T, Moreno-Mayar JV, van Driem G, Gram Wilken U, Seguin-Orlando A, de la Fuente Castro C, Wasef S, Shoocongdej R, Souksavatdy V, Sayavongkhamdy T, Saidin MM, Allentoft ME, Sato T, Malaspinas AS, Aghakhanian FA, Korneliussen T, Prohaska A, Margaryan A, de Barros Damgaard P, Kaewsutthi S, Lertrit P, Nguyen TMH, Hung HC, Minh Tran T, Nghia Truong H, Nguyen GH, Shahidan S, Wiradnyana K, Matsumae H, Shigehara N, Yoneda M, Ishida H, Masuyama T, Yamada Y, Tajima A, Shibata H, Toyoda A, Hanihara T, Nakagome S, Deviese T, Bacon AM, Duringer P, Ponche JL, Shackelford L, Patole-Edoumba E, Nguyen AT, Bellina-Pryce B, Galipaud JC, Kinaston R, Buckley H, Pottier C, Rasmussen S, Higham T, Foley RA, Lahr MM, Orlando L, Sikora M, Phipps ME, Oota H, Higham C, Lambert DM, Willerslev E. 2018. The prehistoric peopling of Southeast Asia. *Science*. 361:88-92.

Nakayama S (Ed.). 2012. Archaeological sites of Gushikawa island: Excavation Report for conservation. *Excavation Report of Okinawa Prefecture Archaeological Center*, Volume 64. (in Japanese).

Nakaza H (Ed.). 2013. Shiraho Saonetabaru Cave Ruins: Emergency Excavation Report due to construction of the New Ishigaki Airport. *Okinawa Prefecture Archaeological Center.* (in Japanese).

Nakaza H (Ed.). 2017a. Shiraho Saonetabaru Cave Ruins: First report on the survey to confirm the extent of important archeological site –Fact report edition–. *Okinawa Prefecture Archaeological Center.* (in Japanese).

Nakaza H (Ed.). 2017b Shiraho Saonetabaru Cave Ruins: Second report on the survey to confirm the extent of important archeological site –Summary report edition–. *Okinawa Prefecture Archaeological Center*. (in Japanese).

Ning C, Li T, Wang K, Zhang F, Li T, Wu X, Gao S, Zhang Q, Zhang H, Hudson MJ, Dong G, Wu S, Fang Y, Liu C, Feng C, Li W, Han T, Li R, Wei J, Zhu Y, Zhou Y, Wang CC, Fan S, Xiong Z, Sun Z, Ye M, Sun L, Wu X, Liang F, Cao Y, Wei X, Zhu H, Zhou H, Krause J, Robbeets M, Jeong C, Cui Y. 2020. Ancient genomes from northern China suggest links between subsistence changes and human migration. *Nat Commun*. 11:2700.

Phillips DL, Koch PL. 2002. Incorporating concentration dependence in stable isotope mixing models. *Oecologia* 130: 114-125.

Raghavan M, Skoglund P, Graf KE, Metspalu M, Albrechtsen A, Moltke I, Rasmussen S, Stafford TW Jr, Orlando L, Metspalu E, Karmin M, Tambets K, Rootsi S, Mägi R, Campos PF, Balanovska E, Balanovsky O, Khusnutdinova E, Litvinov S, Osipova LP, Fedorova SA, Voevoda MI, DeGiorgio M, Sicheritz-Ponten T, Brunak S, Demeshchenko S, Kivisild T, Villems R, Nielsen R, Jakobsson M, Willerslev E. 2013. Upper Palaeolithic Siberian genome reveals dual ancestry of Native Americans. *Nature*. 505:87-91.

Reimer P, Austin W, Bard E, Bayliss A, Blackwell P, Bronk Ramsey C, Butzin M, Cheng H, Edwards R, Friedrich M, Grootes P, Guilderson T, Hajdas I, Heaton T, Hogg A, Hughen K, Kromer B, Manning S, Muscheler R, Palmer J, Pearson C, van der Plicht J, Reimer R, Richards D, Scott E, Southon J, Turney C, Wacker L, Adolphi F, Büntgen U, Capano M, Fahrni S, Fogtmann-Schulz A, Friedrich R, Köhler P, Kudsk S, Miyake F, Olsen J, Reinig F, Sakamoto M, Sookdeo A, Talamo S. 2020. The IntCal20 Northern Hemisphere radiocarbon age calibration curve (0–55 cal kBP). *Radiocarbon* 62: 725-757.

Rohland N, Harney E, Mallick S, Nordenfelt S, Reich D. 2015. Partial uracil-DNA-glycosylase treatment for screening of ancient DNA. *Philos Trans R Soc Lond B Biol Sci.* 370:20130624.

Shimabukuro H (Ed.). 2016. Hanzanbaru A Site. *Chatan Town Board of Education.* (in Japanese).

Shinzato T (Ed.). 2013. Analysis of Tomachin Site at Tokuno Island. *Kagoshima University Archaeological Center.* (in Japanese).

Sakamoto M, Takigami M. 2022. Recalibration of the Radiocarbon Ages of “Yaponesians Genome” Using IntCal20 and Marine20. *Bull. Nat. Mus. Japan*. 237:173-186 (in Japanese).

Shinoda K, Kanzawa-Kiriyama H, Kakuda T, Adachi N. 2019. Genetic characteristics of Yayoi people in Northwestern Kyushu. *Anthropological Sci.* 127:25–43 (in Japanese).

Shinoda K, Kanzawa-Kiriyama H, Kakuda T, Adachi N. 2020. DNA analysis of human bones of the Middle Yayoi period excavated at the Antokudai site, Nakagawa, Fukuoka. *Bull. Nat. Mus. Japan*. 219: 199–206 (in Japanese).

Sikora M, Pitulko VV, Sousa VC, Allentoft ME, Vinner L, Rasmussen S, Margaryan A, de Barros Damgaard P, de la Fuente C, Renaud G, Yang MA, Fu Q, Dupanloup I, Giampoudakis K, Nogués-Bravo D, Rahbek C, Kroonen G, Peyrot M, McColl H, Vasilyev SV, Veselovskaya E, Gerasimova M, Pavlova EY, Chasnyk VG, Nikolskiy PA, Gromov AV, Khartanovich VI, Moiseyev V, Grebenyuk PS, Fedorchenko AY, Lebedintsev AI, Slobodin SB, Malyarchuk BA, Martiniano R, Meldgaard M, Arppe L, Palo JU, Sundell T, Mannermaa K, Putkonen M, Alexandersen V, Primeau C, Baimukhanov N, Malhi RS, Sjögren KG, Kristiansen K, Wessman A, Sajantila A, Lahr MM, Durbin R, Nielsen R, Meltzer DJ, Excoffier L, Willerslev E. 2019. The population history of northeastern Siberia since the Pleistocene. *Nature.* 570:182-188.

Stuiver M, Reimer PJ. 1993. Extended ^14^C data base and revised CALIB 3.0 ^14^C age calibration program. *Radiocarbon*. 35: 215-230.

Takamiya H (Ed.). 1993. Excavation survey summary of Ufutobaru Shell Mounds at Yomitan Village. *Bulletin of Yomitan Village History and Folklore Museum.* 17:1-47. (in Japanese).

Takenaka M. 2013. Human bones excavated from Tomachin Site. In Shinzato T (Ed.), Analysis of Tomachin Site at Tokuno Island (pp. 149-162). *Kagoshima University Archaeological Center.* (in Japanese).

Takenaka M, Sakamoto M, Takigami M. 2021. Archaeological Report on the Chronology of Human Bones Excavated in Tokunoshima Island, Kagoshima Pref. : Omonawa No.1 Shell Midden, Tomachin Site and Shitabaru Cave Site. *Bull. Nat. Mus. Japan*. 228:441-448 (in Japanese).

Tokumine R, Katagiri C, Kobashigawa T, Motomura M, Oshiro A, Tengan M, Sugawara H, Doi N, Yoneda M. 2009. Human Remains recovered from the Shiru area of Furuzamamibaru district, Zamami Village, Okinawa Prefecture. *Bulletin of the Archaeological study of Okinawa.* 6:41-52 (in Japanese).

Toma T. 1975. Emergency Excavation Report of Tsuken Second Shell Mounds: Flexed burial remains. In Annual Report of Cultural Properties for Fiscal Year Showa 49 (pp.64). *Okinawa Prefectural Board of Education*. (in Japanese).

Tomoyori E. 1968. Summary of Gushibaru Shell Mound at Ie island. *Bulletin of the College of Law and Letters, University of the Ryukyus. Sociology*, volume 11. (in Japanese).

van Klinken GL. 1999. Bone collagen Quality Indicators for palaeodietary and radiocarbon measurements. *J. Archaeol. Sci.* 26: 687-695.

Yang MA, Gao X, Theunert C, Tong H, Aximu-Petri A, Nickel B, Slatkin M, Meyer M, Pääbo S, Kelso J, Fu Q. 2017. 40,000-Year-Old Individual from Asia Provides Insight into Early Population Structure in Eurasia. *Curr Biol.* 27:3202-3208.e9.

Yang MA, Fan X, Sun B, Chen C, Lang J, Ko YC, Tsang CH, Chiu H, Wang T, Bao Q, Wu X, Hajdinjak M, Ko AM, Ding M, Cao P, Yang R, Liu F, Nickel B, Dai Q, Feng X, Zhang L, Sun C, Ning C, Zeng W, Zhao Y, Zhang M, Gao X, Cui Y, Reich D, Stoneking M, Fu Q. 2020. Ancient DNA indicates human population shifts and admixture in northern and southern China. *Science*. 369:282-288.

Yoneda M, Uno H, Shibata Y, Suzuki R, Kumamoto Y, Yoshida K, Sasaki T, Suzuki A, Kawahata H. 2007. Radiocarbon marine reservoir ages in the western Pacific estimated by pre-bomb molluscan shells. *Nuclear Instruments and Methods in Physics Research B*. 259:432-437.

Wang CC, Yeh HY, Popov AN, Zhang HQ, Matsumura H, Sirak K, Cheronet O, Kovalev A, Rohland N, Kim AM, Mallick S, Bernardos R, Tumen D, Zhao J, Liu YC, Liu JY, Mah M, Wang K, Zhang Z, Adamski N, Broomandkhoshbacht N, Callan K, Candilio F, Carlson KSD, Culleton BJ, Eccles L, Freilich S, Keating D, Lawson AM, Mandl K, Michel M, Oppenheimer J, Özdoğan KT, Stewardson K, Wen S, Yan S, Zalzala F, Chuang R, Huang CJ, Looh H, Shiung CC, Nikitin YG, Tabarev AV, Tishkin AA, Lin S, Sun ZY, Wu XM, Yang TL, Hu X, Chen L, Du H, Bayarsaikhan J, Mijiddorj E, Erdenebaatar D, Iderkhangai TO, Myagmar E, Kanzawa-Kiriyama H, Nishino M, Shinoda KI, Shubina OA, Guo J, Cai W, Deng Q, Kang L, Li D, Li D, Lin R, Nini, Shrestha R, Wang LX, Wei L, Xie G, Yao H, Zhang M, He G, Yang X, Hu R, Robbeets M, Schiffels S, Kennett DJ, Jin L, Li H, Krause J, Pinhasi R, Reich D. 2021. Genomic insights into the formation of human populations in East Asia. Nature. 591:413-419.

Zamami Village History Editing Committee. 1989. Summary of in-village site. In History of Zamami Village, Volume 1 (pp. 101-123). *Zamami Village History Editing Committee.* (in Japanese).
